# Probabilistic Sensory Constraints Shape Swarm Cohesion in Echolocating Bats

**DOI:** 10.1101/2025.07.05.663265

**Authors:** Ravi Umadi

**Author notes:** Corresponding author:* Ravi Umadi–.

## Abstract

1. Cohesive animal groups rely on continuous behavioural updating to regulate spacing, alignment, and collision avoidance, yet all sensory-guided interaction is constrained by finite signal propagation delays, processing time, and motor latency. In actively sensing species such as echolocating bats, these constraints raise a fundamental ecological question: *what limits the stability and density of cohesive groups under increasing interaction load?*
2. I develop a constraint-based framework in which neighbour-based interaction is treated as a closed-loop process that is only feasible when sensory updates can be acquired, processed, and acted upon within a finite temporal budget. Building on an asynchronous swarm simulation grounded in echo-timed biosonar control, I formalise two receiver-side feasibility constraints that become critical in dense groups: (i) temporal overlap between conspecific calls and the echo-processing window, and (ii) level dominance of the tracked neighbour’s echo over competing conspecific signals.
3. Both constraints emerge as *probabilistic feasibility boundaries* rather than binary conditions for perceptual success or failure. Across a broad parameter space of responsivity, group density, and flight speed, the fraction of call events supporting reliable neighbour-based updates declines smoothly with compounded interaction load. Temporal overlap is organised by a single compound term integrating local density, neighbourhood call rate, call duration, and echo delay, while level dominance operates in a marginal regime where masking is frequent but intermittent.
4. Simulation results show that stable, collision-averse swarm cohesion can persist across wide ranges of density and motion without explicit coordination or interference avoidance, provided that sufficiently frequent informative updates remain available. Breakdown occurs not because echoes become undetectable, but because the probability of obtaining timely, behaviourally relevant updates falls below what is required to sustain closed-loop control.
5. Together, these findings identify temporal feasibility and interaction dominance as general constraints shaping cohesion, spacing, and fragmentation in actively sensing animal groups. Rather than invoking acoustic jamming as a failure mode, the framework reframes interference as a background condition that regulates the statistics of actionable information. This closed-loop perspective provides a unifying ecological principle for collective behaviour in active sensing systems and generates testable predictions for when cohesion should persist or fail under increasing sensory and interactional demand.

## 1 INTRODUCTION

Collective motion in animals emerges from local interactions that are constrained by the sensory, biomechanical, and cognitive capacities of individuals. Across taxa, cohesive groups must reconcile the benefits of aggregation—such as predator avoidance, information sharing, and navigational efficiency—with the risks of collision, interference, and information overload [1–3]. A central challenge in animal ecology is therefore to understand how simple local rules remain viable under increasing density, and which constraints ultimately limit the scale and structure of collective behaviour.

Echolocating bats provide an extreme and informative test case for this problem. During emergence from large roosts, thousands of individuals fly in close proximity while relying on self-generated acoustic signals for navigation and collision avoidance [4–7]. Each bat must emit calls, receive echoes, process acoustic information, and update flight trajectories within tens of milliseconds, all while surrounded by intense conspecific acoustic activity. This situation has long been framed as an acoustic “cocktail-party problem” [8], in which overlapping calls and echoes threaten the reliable interpretation of self-generated sensory information [9–11].

A substantial body of work has examined how bats cope with acoustic interference in groups. Experimental and modelling studies have shown that bats can often detect relevant echoes despite severe masking, and that they adjust call parameters such as intensity, duration, and rate in response to conspecific noise [12, 13]. Agent-based simulations incorporating auditory receivers suggest that some strategies, such as spectral jamming avoidance, may offer limited benefit under dense social conditions [14]. More recently, field studies combining onboard microphones and trajectory reconstruction have demonstrated that bats can maintain low collision risk during emergence even when a large fraction of echoes are masked, largely through rapid spatial spreading after leaving the roost [15]. Together, these studies establish that bats are remarkably robust to interference—but they leave open a deeper question: *what ultimately constrains the feasibility of neighbour-based interactions as density increases?*

Most existing approaches treat masking and jamming as *outcomes* of prescribed behaviours. Call parameters and interaction rules are assumed, and the resulting detectability, jamming probability, or collision rate is measured. While this has yielded important insights, it does not address whether a given interaction rule can remain dynamically self-consistent under crowding. In particular, echolocation is an intrinsically closed-loop process: the timing of call emission, echo reception, neural processing, and motor response are tightly coupled, and delays in this loop directly limit how rapidly and accurately an animal can respond to changes in its environment [16, 17]. When multiple individuals rely on similar closed loops while interacting in space, collective behaviour is not only shaped by perception, but bounded by the temporal feasibility of those loops.

Here, I develop a mechanistic framework in which swarm cohesion emerges from *stable temporal loops* linking inter-individual distance to echo delay, call timing, and behavioural response. Building on a nearest-neighbour interaction model, I show that closed-loop timing alone implies a lower bound on the temporal budget available for collision avoidance and cohesion. I then extend this framework by explicitly incorporating two receiver-side constraints that become critical in dense groups: *temporal overlap* and *level dominance*. Temporal overlap captures the probability that conspecific call onsets intrude into the time window required to interpret self-generated echoes, while level dominance formalises the requirement that the echo from the next neighbour exceeds competing conspecific signals by a biologically meaningful margin.

Crucially, these constraints are not imposed heuristically. Temporal overlap is derived from the statistics of conspecific call arrivals within an attenuation- and directionality-limited interaction volume, yielding an explicit density-dependent bound on viable call timing and duration. Level dominance is derived from acoustic attenuation and stochastic geometry, linking source level, neighbour distance, and swarm density to the probability that the nearest neighbour’s echo remains salient. Together, these constraints transform masking from a descriptive phenomenon into a set of *feasibility conditions* that bound which combinations of call rate, call duration, and call level can sustain neighbour-based cohesion.

This perspective reframes several empirical observations. Adjustments such as calling louder, longer, or more frequently can be interpreted as attempts to satisfy competing constraints, but they need not reduce interference and may even exacerbate it under high density. Spatial spreading during emergence emerges not merely as a behavioural choice, but as a means of restoring temporal and acoustic feasibility. Moreover, the framework predicts density-dependent regimes in which self-echo-based neighbour tracking becomes untenable, motivating transitions to alternative interaction strategies, such as coarser neighbour selection or soundscape-based navigation [18].

By integrating closed-loop timing with explicit overlap and dominance constraints, this study provides a general, analytically tractable account of how sensory limitations shape collective behaviour. Although developed in the context of echolocating bats, the framework addresses a broader ecological problem: how local, delay-limited sensing can sustain—or fail to sustain—cohesion in dense animal groups. The results generate testable predictions about call timing, source levels, and spacing during emergence, and clarify the conditions under which behavioural transitions should occur as density increases.

## 2 METHODS

### 2.1 Overview

I developed an asynchronous, event-driven simulation to test whether purely local, sonar-guided control rules can support stable, cohesive, and collision-averse flight in an emerging bat swarm. The model is grounded in the responsivity framework [19, 20] and extends it to multi-agent flight by coupling each individual’s locomotor updates to an echo-timed sensorimotor loop. In addition to the kinematic and temporal backbone that generates cohesion, I introduce two receiver-side feasibility constraints motivated by the acoustic “cocktail party” problem in dense groups: (i) *temporal overlap*, quantifying the probability that conspecific signals intrude into the focal bat’s echo-processing window; and (ii) *level dominance*, requiring that the focal bat’s echo from its next neighbour remains reliably more salient than competing conspecific signals. Together, the simulation and the analytical constraints provide a predictive basis for density-dependent call adjustments and for regime shifts in which neighbour-based echo tracking becomes unreliable.

All simulations were implemented in Matlab (R2023b). The full history of each bat’s state and call timing was logged for quantitative analysis.

### 2.2 Swarm simulation model and asynchronous implementation

I simulate a swarm of *N* bats (agents) indexed by *i* ∈ {1, …, *N*} with continuous-time positions **p**_*i*_(*t*) ∈ ℝ^3^, unit heading vectors **h**_*i*_(*t*), and speeds *v*_*i*_(*t*). Each bat operates an independent sonar-control loop with its own call schedule. The simulation is *asynchronous*: at any event time, only one bat emits a call and updates its sensory estimate and motor state, while all bats continue to move continuously between events.

#### Initialisation

Initial positions are sampled from a three-dimensional Gaussian distribution with user-defined spread *σ*_init_ (standard deviation per axis),

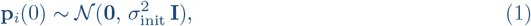

where **I** is the 3 × 3 identity matrix. Initial headings are aligned with the +*X* axis with small angular perturbations, and initial speeds are sampled from a narrow uniform range (±10%) around a nominal target speed *v*_0_.

#### Event-driven stepping

Let 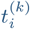 be the scheduled time of bat *i*’s *k*th call. The simulation proceeds by selecting the minimum scheduled event time,

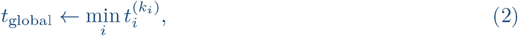

advancing the global time to *t*_global_, and forward-integrating all positions from their last update times to *t*_global_. The calling bat then executes: (i) target selection and echo-timed update of call timing; (ii) heading and speed update; and (iii) rescheduling of its next call time by adding its current interpulse interval. (see pseudocode in Algorithm A.1.)

### 2.3 Sonar timing backbone: echo delay, responsivity, IPI, and call duration

At each call iteration, bat *i* selects a target distance *d*_*i*_(*t*) according to a simple frontal-selection rule:

- If at least one neighbour lies ahead (defined by a positive dot product between the focal heading **h**_*i*_ and the displacement to that neighbour), *d*_*i*_(*t*) is set to the Euclidean distance to the nearest forward neighbour.
- If no bat is detected ahead, a virtual obstacle is assigned at a random distance *d*_obs_ ∈ [2, 5] m to maintain a biologically plausible baseline call rate in open space.

The round-trip echo delay is

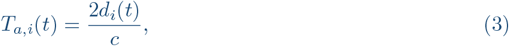

where *c* = 343 m s^−1^ is the speed of sound. The behavioural response interval (the time budget between informative echo receipt and enacting a motor update) is represented as a proportional margin on echo delay:

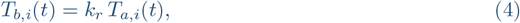

where *k*_*r*_ *>* 0 is a dimensionless *responsivity coefficient*. The resulting interpulse interval (IPI) is

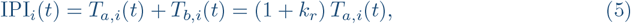

with an enforced maximum call rate of 200 Hz.

#### Call duration

Call duration is adapted in an echo-timed manner with a biologically motivated lower bound. Let *T*_call,*i*_(*t*) denote the call duration (s), with *T*_call,min_ = 0.5 ms and baseline *T*_call,init_. I implement a contraction rule:

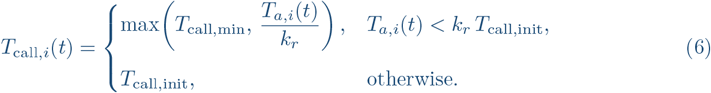

For analytical developments below, I denote call duration by *τ* (*t*) and allow a more general contraction factor *κ* such that *τ* = clip(*T*_*a*_*/κ*; *τ*_min_, *τ*_max_); the special case *κ* = *k*_*r*_ recovers the simulation rule in Eq. (6).

### 2.4 Flight dynamics

Each bat’s position is updated continuously as:

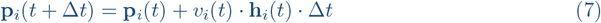

where **p**_*i*_(*t*) is the 3D position of bat *i* at time *t, v*_*i*_(*t*) its current velocity (speed), **h**_*i*_(*t*) its unit heading vector (direction of flight), and Δ*t* is the time elapsed since the last update of that bat’s position. In the asynchronous simulation, Δ*t* differs for each bat and corresponds to its current inter-pulse interval (IPI), thus coupling locomotor updates to sonar control.

Heading and velocity are updated only when each bat emits a sonar call, i.e. at its own echo-processing iteration. At each such iteration, the heading vector **h**_*i*_ is adjusted by a biologically inspired rule combining several components:

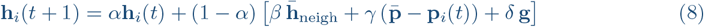

where:

- *α* = 0.9 is an inertia term promoting smooth directional changes.
- *β, γ, δ* are weighting parameters for the three control components:
  - **Neighbour alignment** (*β*): tendency to align heading with the mean heading of the other bats, 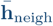.
  - **Cohesion to swarm centroid** (*γ*): damping pull toward the average position of the swarm, 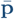.
  - **Global drive** (*δ*): bias toward an overall goal direction **g** (set along +*X*).

This update rule belongs to the general class of alignment–cohesion–drive models widely used to describe collective animal behaviour [3, 21]. It was specifically adapted here to reflect the biological constraints of bat flight, which requires maintaining forward progression while preserving local cohesion.

Individual velocities *v*_*i*_(*t*) are also allowed to drift gradually between [0.8, 1.2]×*v*_0_, introducing naturalistic variation in speed between individuals. Velocity changes occur stochastically, with small random increments drawn at each bat’s call iteration.

Through this scheme, both positional updates and directional dynamics are controlled asynchronously and locally, with each bat responding only to its own sensory input (echoes) and the perceived state of nearby conspecifics.

Speeds are allowed to drift gradually within [0.8, 1.2] *v*_0_ via small stochastic increments applied at call events, introducing heterogeneity consistent with natural variation.

### 2.5 Collision risk detection

To evaluate the swarm’s ability to maintain safe spacing, I implemented a collision risk detection module based on two criteria.

First, a critical response threshold 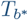 was defined, representing the minimum physiological time required for bats to react to a perceived obstacle. Following the responsivity framework of [19], I used 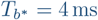 ms. If, at any sonar iteration:

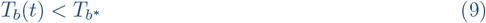

a collision risk is flagged for the emitting bat.

Second, a time-to-collision (TTC) metric was computed at each global iteration for every pair of bats (*m, n*), based on their current relative position and velocity. For a pair of bats separated by distance *d* and relative closing velocity *v*_closing_:

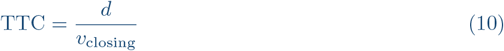

if *v*_closing_ *>* 0; otherwise, TTC = ∞.

A collision risk was logged if:

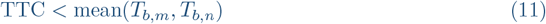

where *T*_*b,m*_ and *T*_*b,n*_ are the current echo-response intervals of the two bats. To avoid redundant reporting, a cooldown interval of 0.1 s was enforced between successive alerts for the same pair.

These criteria allow explicit tracking of instances where bats’ current sonar update timing would no longer suffice to avoid imminent collisions, providing a quantitative measure of the swarm’s collective safety margin.

### 2.6 Receiver-side constraint criteria: temporal overlap and level dominance

The swarm simulation above assumes that, at each call event, a bat can accommodate the receiver-side processing demands required to extract self-generated echoes from its selected neighbour. In dense groups, this assumption can break down in a probabilistic manner: conspecific calls can temporally overlap the echo-processing window, and competing direct calls can exceed the echo level at the receiver. I therefore formalise these effects as receiver-side *constraint criteria* that quantify the risk of temporal overlap and level-dominance violation.

#### 2.6.1 Temporal overlap constraint

Let *ρ* denote local swarm number density (bats m^−3^). Only a subset of conspecifics contribute substantial interference, determined by directional geometry and attenuation. I represent this as an effective interference volume

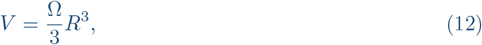

where Ω is the effective solid angle (sr) of relevant interference (e.g. a forward sector) and *R* is an interference radius (m) defined by audibility or masking thresholds (below).

Let 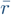 be the mean call rate (Hz) of potential interferers. The aggregate arrival rate of conspecific call onsets at the focal receiver is approximated as

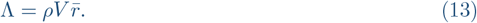

I define a vulnerable window *W* (s) as the interval in which interference is most likely to disrupt echo interpretation:

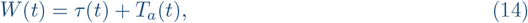

where *τ* (*t*) is call duration and *T*_*a*_(*t*) is echo delay (Eq. (3)). Approximating conspecific call onsets as a Poisson point process at the receiver—a standard assumption for stochastic interference modelling in communication networks [22–24], the probability of at least one overlap event is

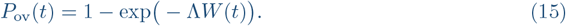

I impose a tolerable overlap constraint

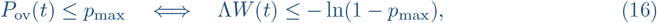

which yields an explicit density–timing inequality:

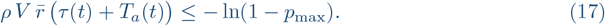

For analysis and visualisation, I express temporal overlap in terms of a dimensionless boundary ratio,

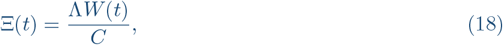

where *C* = − ln(1 − *p*_max_) is the maximum tolerable information load implied by the overlap tolerance *p*_max_. Values Ξ *<* 1 indicate operation within the receiver-side feasibility region, whereas Ξ *>* 1 indicate that the expected temporal overlap probability exceeds the tolerable bound. Importantly, Ξ does not impose a hard constraint on behaviour; it serves as a normalised diagnostic of proximity to the temporal feasibility boundary and is used for run-level comparison across parameter regimes.

#### 2.6.2 Acoustic attenuation and interference radius

Let *SL* denote source level (dB re 20 *µ*Pa at 1 m) and *f* a dominant frequency. One-way transmission loss is modelled as

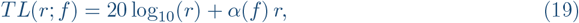

where *α*(*f*) is atmospheric absorption (dB m^−1^). The received level of a direct conspecific call at range *r* is

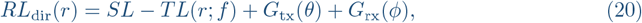

where *G*_tx_ and *G*_rx_ are directional gains (dB). The interference radius *R* is defined implicitly by an audibility/masking threshold Θ_dir_:

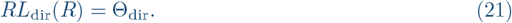

In the isotropic case (*G*_tx_ = *G*_rx_ = 0), this becomes

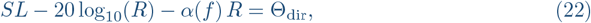

which is solved numerically to obtain *R*(*SL, f*, Θ_dir_) and hence *V* (*SL, f*) in Eq. (12).

#### 2.6.3 Level dominance constraint

Let *TS* denote an effective target strength (dB) for the tracked neighbour. The received echo level from the neighbour at distance *d* is

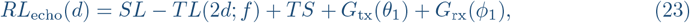

where propagation distance is 2*d* (round-trip). Let *RL*_max_ be the level of the strongest interfering direct call among interferers:

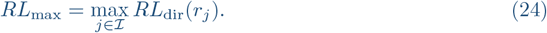

I require a salience margin Δ (dB) such that

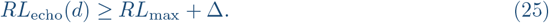

To express this in density-dependent form, I approximate the spatial configuration of interferers in the sector as a Poisson point process with intensity *ρ*. The survival function of the nearest interferer distance *r*_min_ is

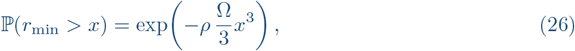

yielding the *q*-quantile

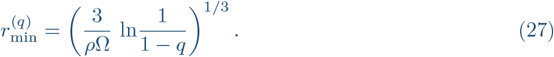

Under a conservative approximation that the nearest interferer dominates,

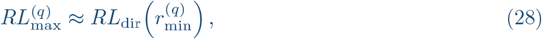

and the probabilistic dominance requirement becomes

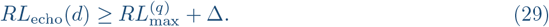

The quantile parameter *q* controls how conservatively dominance is enforced, specifying the fraction of call events for which the tracked neighbour’s echo is expected to exceed competing calls by the required margin Δ.

#### 2.6.4 Joint constraints and predicted adjustments

I treat temporal overlap and level dominance as simultaneous constraints defining an accommodative region for call duration, call rate, and source level:

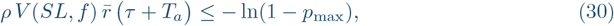

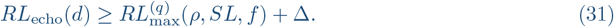

Because increasing *SL* increases both echo level and interference radius (hence *V*), these constraints naturally generate trade-offs and, in crowded regimes, density-dependent optima for *SL* and *τ*.

### 2.7 Parameter sweeps and simulation outputs

For each simulation run, I recorded full time series for each bat: positions **p**_*i*_(*t*), headings **h**_*i*_(*t*), speeds *v*_*i*_(*t*), target distances *d*_*i*_(*t*), echo delays *T*_*a,i*_(*t*), response intervals *T*_*b,i*_(*t*), IPI IPI_*i*_(*t*), call durations *T*_call,*i*_(*t*), neighbour identities, and collision-risk events (Eqs. (7)–(11)). These histories were used to compute summary statistics of inter-individual spacing, call timing distributions, and collision-risk rates.

To assess how responsivity and speed jointly shape swarm dynamics, I conducted grid simulations varying *k*_*r*_ and *v*_0_ across predefined sets (Tables 1 and 2). Each parameter combination was simulated for 60 s with *N* = 50 bats. An extended set of simulations with *σ*_*init*_ = [1, 2, 3] (see code [25]) was run to evaluate the group density effects. Runs were executed in parallel (`parfor`), and I quantified total collision-risk events, normalised collision rates, and descriptive statistics (mean, median, standard deviation) for call duration, call rate, velocity, and neighbour distance.

**Table 1:** Summary statistics of swarm simulations across different combinations of initial velocity *v*_0_ and responsivity coefficient *k*_*r*_. For each simulation, the table reports the mean (*µ*), median (med.), and standard deviation (*σ*) of call duration (CD), call rate (CR), velocity (*v*), and inter-bat distance (*d*), averaged over all individuals and time points.

| $k_r$ | $v_0$ | CD($\mu$) | CD(med.) | CD($\sigma$) | CR($\mu$) | CR(med.) | CR($\sigma$) | $v(\mu)$ | $v(\text{med.})$ | $v(\sigma)$ | $d(\mu)$ | $d(\text{med.})$ | $d(\sigma)$ |
| --- | --- | --- | --- | --- | --- | --- | --- | --- | --- | --- | --- | --- | --- |
| 3.0 | 3.0 | 0.930 | 0.500 | 1.2 | 151.2 | 196.5 | 61.1 | 3.0 | 3.0 | 0.348 | 6.4 | 5.7 | 3.7 |
| 3.0 | 4.0 | 1.1 | 0.500 | 1.5 | 143.7 | 179.8 | 65.2 | 3.9 | 3.9 | 0.427 | 5.1 | 4.0 | 4.7 |
| 3.0 | 5.0 | 1.7 | 0.653 | 2.3 | 122.2 | 127.7 | 71.4 | 4.9 | 4.9 | 0.535 | 7.3 | 5.0 | 9.6 |
| 3.0 | 6.0 | 1.7 | 0.676 | 2.3 | 120.9 | 123.3 | 72.6 | 5.8 | 5.8 | 0.562 | 9.9 | 6.6 | 12.6 |
| 3.0 | 7.0 | 1.7 | 0.640 | 2.3 | 122.0 | 130.1 | 72.4 | 6.8 | 6.8 | 0.648 | 7.0 | 4.6 | 7.2 |
| 4.0 | 3.0 | 0.811 | 0.500 | 0.973 | 136.7 | 153.7 | 64.9 | 3.0 | 2.9 | 0.343 | 6.3 | 5.8 | 3.6 |
| 4.0 | 4.0 | 1.1 | 0.500 | 1.5 | 116.2 | 112.9 | 70.0 | 3.9 | 3.8 | 0.458 | 11.3 | 10.1 | 6.8 |
| 4.0 | 5.0 | 1.3 | 0.500 | 1.7 | 116.0 | 117.1 | 72.8 | 5.0 | 5.0 | 0.526 | 7.3 | 4.7 | 8.0 |
| 4.0 | 6.0 | 1.3 | 0.500 | 1.8 | 117.8 | 117.0 | 72.0 | 5.9 | 5.9 | 0.611 | 8.7 | 5.5 | 10.9 |
| 4.0 | 7.0 | 1.4 | 0.539 | 1.9 | 106.9 | 92.8 | 72.4 | 6.8 | 6.8 | 0.661 | 8.1 | 4.7 | 10.9 |
| 5.0 | 3.0 | 0.700 | 0.500 | 0.809 | 138.7 | 163.3 | 66.4 | 2.9 | 2.9 | 0.332 | 5.1 | 4.1 | 4.5 |
| 5.0 | 4.0 | 0.878 | 0.500 | 1.1 | 115.5 | 109.8 | 70.5 | 3.9 | 3.9 | 0.439 | 10.2 | 5.4 | 10.0 |
| 5.0 | 5.0 | 1.0 | 0.500 | 1.4 | 111.0 | 104.6 | 71.7 | 4.7 | 4.6 | 0.498 | 14.3 | 6.9 | 18.6 |
| 5.0 | 6.0 | 0.997 | 0.500 | 1.5 | 112.3 | 104.5 | 71.4 | 5.8 | 5.7 | 0.533 | 8.8 | 5.3 | 12.0 |
| 5.0 | 7.0 | 1.2 | 0.500 | 1.7 | 103.4 | 92.3 | 71.1 | 6.8 | 6.8 | 0.564 | 10.6 | 7.8 | 12.5 |
| 6.0 | 3.0 | 0.681 | 0.500 | 0.781 | 126.0 | 131.2 | 68.2 | 3.0 | 2.9 | 0.348 | 6.9 | 4.3 | 7.3 |
| 6.0 | 4.0 | 0.741 | 0.500 | 0.942 | 118.5 | 116.5 | 68.8 | 3.9 | 3.9 | 0.422 | 5.4 | 3.9 | 5.2 |
| 6.0 | 5.0 | 0.927 | 0.500 | 1.3 | 112.2 | 108.6 | 73.5 | 5.0 | 5.0 | 0.514 | 14.3 | 11.2 | 11.1 |
| 6.0 | 6.0 | 0.944 | 0.500 | 1.3 | 92.7 | 73.1 | 68.6 | 5.8 | 5.8 | 0.540 | 12.0 | 8.2 | 10.7 |
| 6.0 | 7.0 | 1.1 | 0.500 | 1.5 | 85.9 | 64.0 | 69.0 | 6.8 | 6.8 | 0.550 | 9.6 | 5.7 | 11.2 |
| 7.0 | 3.0 | 0.656 | 0.500 | 0.676 | 107.7 | 97.3 | 68.3 | 2.9 | 2.9 | 0.355 | 5.5 | 4.1 | 5.1 |
| 7.0 | 4.0 | 0.727 | 0.500 | 0.882 | 111.3 | 102.9 | 71.8 | 3.9 | 3.9 | 0.404 | 7.0 | 4.6 | 6.8 |
| 7.0 | 5.0 | 0.814 | 0.500 | 1.1 | 93.7 | 74.8 | 69.6 | 4.7 | 4.6 | 0.487 | 10.7 | 6.2 | 13.4 |
| 7.0 | 6.0 | 0.838 | 0.500 | 1.2 | 96.2 | 77.4 | 70.3 | 5.8 | 5.8 | 0.497 | 10.7 | 5.4 | 13.6 |
| 7.0 | 7.0 | 0.951 | 0.500 | 1.3 | 81.8 | 51.5 | 70.5 | 6.7 | 6.6 | 0.589 | 17.3 | 12.6 | 16.4 |

**Table 2:**
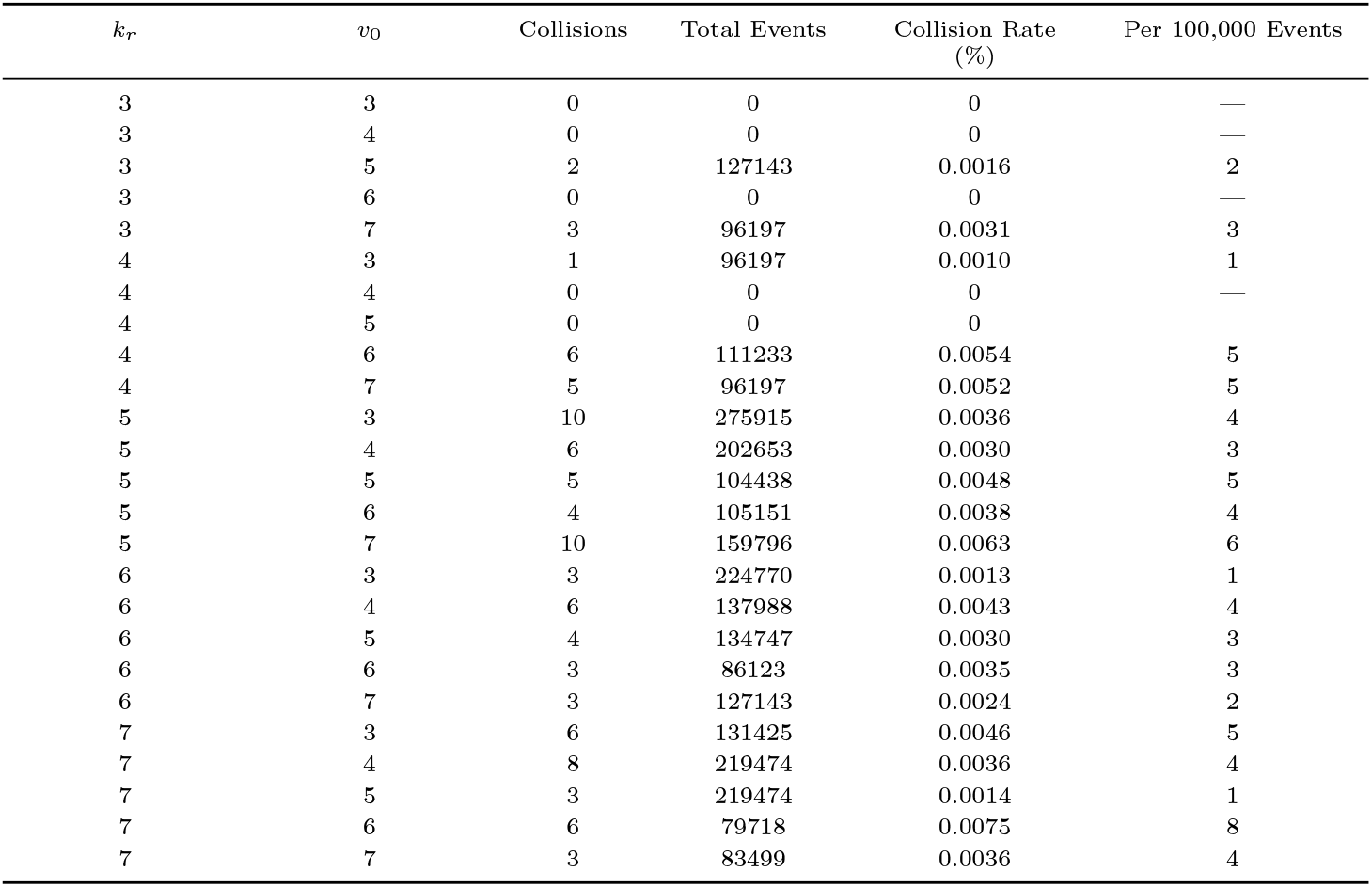
Summary of collision events across simulation runs for different combinations of responsivity coefficient *k*_*r*_ and initial velocity *v*_0_. The table reports the absolute number of collisions, total simulated events, collision rate as a percentage, and normalised rate per 100,000 call events. Collision rates remain low overall, but increase slightly with higher *k*_*r*_ and *v*_0_ values, suggesting that slower responsivity (higher *k*_*r*_) or higher velocities may modestly elevate collision risk within the swarm.

### 2.8 Modelling information transfer in asynchronous biosonar swarms

To quantify how a local perturbation propagates through a neighbour-coupled, echo-timed swarm, I derived a minimal model of spatiotemporal information flow along a chain of *n* interactions.

#### 2.8.1 Information latency and information velocity

Let *d*_*i*_ denote the distance between bat *i* and its nearest frontal neighbour. The corresponding echo delay is *T*_*a,i*_ = 2*d*_*i*_*/c*. Each bat reacts after a latency *T*_*b,i*_ = *k*_*r*_*T*_*a,i*_, governed by the responsivity coefficient *k*_*r*_. As a perturbation propagates across *n* agents, the total distance covered is 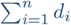, and the cumulative delay is 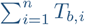.

I define the following two quantities:

- **Information latency** *I*_*t*_(*n*): the time taken for a perturbation to reach the *n*-th agent in the chain.
- **Information velocity** *I*_*v*_: the effective speed at which a perturbation propagates across the swarm.

The information latency is:

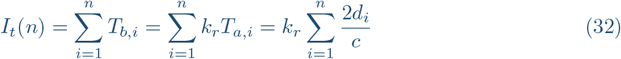

The effective information velocity is then given by:

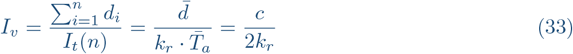

where 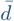 is the mean inter-bat distance, and 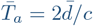 is the average echo delay.

This formulation accounts for spatial heterogeneity and reveals how *I*_*v*_ depends primarily on the biosonar responsivity *k*_*r*_ and the speed of sound *c*, while *I*_*t*_(*n*) depends on both *k*_*r*_ and the summed echo delays across *n* agents. Denser regions (smaller 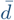) yield shorter echo delays and smaller *I*_*t*_(*n*), facilitating faster local corrections. In contrast, sparser regions increase both delay and spatial reach per step, but the overall information velocity *I*_*v*_ remains invariant for constant *k*_*r*_. Thus, *k*_*r*_ regulates the trade-off between speed and stability: lower values enhance corrective responsiveness, while higher values promote robustness by damping rapid fluctuations.

#### 2.8.2 Perturbation Attenuation Model

As a perturbation propagates through the swarm, its influence may diminish due to accumulated noise, variability in heading updates, or stochastic deviations in response. To model this attenuation, I assume that each bat transmits a fraction of the received perturbation to its neighbour, resulting in a multiplicative loss over distance. This leads to an exponential decay of the form:

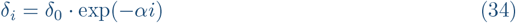

where:

- *δ*_*i*_ is the perturbation magnitude perceived by the *i*^th^ bat in the interaction chain,
- *δ*_0_ is the initial perturbation magnitude,
- *α* is the decay coefficient, incorporating the effect of divergence, local noise, and behavioural variability.

To estimate how far a perturbation spreads before becoming negligible, I define a behavioural threshold *δ*_min_ below which the perturbation has no effective influence. The maximum range of influence (in terms of hops) is then:

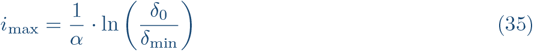

This defines the *effective communication range* of a perturbation within the swarm. For a given *α*, smaller initial perturbations or stricter behavioural thresholds reduce the range of effect, while lower *α* (i.e., more consistent responses) extend the range.

My modelling framework aligns closely with the principles outlined in the general theory of information transfer by Ahlswede (2008) [26], particularly in its treatment of information as a spatiotemporal dynamic process shaped by system structure and transmission constraints. In his formalism, information transfer is not merely a passive relay but an active transformation governed by local processing rules and communication delays. Similarly, I define information flow through a biologically grounded mechanism: each agent in the swarm reacts based on temporally precise biosonar feedback, creating a sequential chain of responses. This structure directly mirrors the discrete, causally ordered channels in Ahlswede’s theory. Moreover, my explicit formulation of information velocity and perturbation attenuation offers a quantitative instantiation of the abstract transfer functions, adapted for real-time, distributed sensory systems such as bat swarms. In both frameworks, the fidelity and speed of information propagation are critically dependent on the internal system dynamics—in the current case, the responsivity coefficient *k*_*r*_ and the spatial heterogeneity of the swarm.

#### 2.8.3 Interpretation and Application

This model provides a quantitative framework for analysing the swarm’s sensitivity to local disturbances. It shows that:

- The reaction time (*T*_*b*_), governed by *k*_*r*_, sets the temporal resolution of feedback within the swarm.
- Dense swarms (small *d*) transmit information faster, but also require more frequent updates to maintain coherence.
- There exists a trade-off: higher responsivity improves reaction speed but reduces tolerance to noise, while lower responsivity improves robustness at the cost of slower adaptation.

In my grid-based simulation, each point in the parameter space defined by (*k*_*r*_, *v*_0_) corresponds to a different regime of this trade-off. The information velocity model thus allows theoretical prediction of:

1. Whether a perturbation will propagate fast enough to be corrected
2. How far it will spread before becoming negligible
3. The role of *k*_*r*_ in mediating this behaviour across the swarm

This model provides a theoretical lens through which simulation outcomes—such as inter-bat distances, collision rates, and velocity fluctuations—can be interpreted as emergent consequences of biosonar loop dynamics and local sensory delays.

### 2.9 Receiver-side constraint criteria in simulation

The receiver-side constraint criteria (Section 2.6) are used in *post-hoc* receiver-side constraint analysis, within the simulation framework.

For each call event of bat *i*, the simulation provides the realised interaction distance *d*_*i*_(*t*), echo delay *T*_*a,i*_(*t*), call duration *T*_call,*i*_(*t*) (Eq. (6)), and call rate *r*_*i*_(*t*) = IPI_*i*_(*t*)^−1^ (Eq. (5)). Given a local density estimate *ρ*_*i*_(*t*) (defined below) and chosen acoustic parameters (*SL, f*, Θ_dir_, *TS*, Δ), I evaluate two receiver-side constraint criteria that quantify the *probabilistic risk* of echo interference.

First, the probability that conspecific calls overlap the effective echo-processing window is evaluated using the compound load term Λ*W*, such that

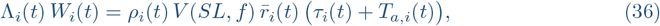

where 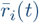 is the local neighbourhood call rate and *W*_*i*_(*t*) is the effective temporal window. This quantity is mapped to an overlap probability

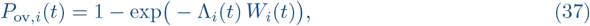

which is compared against a reference bound *p*_max_ via the criterion

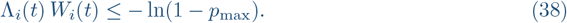

Second, level dominance is evaluated by comparing the received echo level from the tracked neighbour with the *q*-quantile of the expected distribution of direct-call levels from surrounding bats:

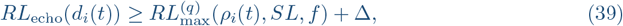

where Δ is a conservative dominance margin.

Neither criterion is imposed as a hard constraint during simulation. Instead, each call event is assigned a time-resolved indicator reflecting whether it falls within a region of low expected overlap probability and positive dominance margin. These indicators are subsequently summarised at the run level as proportions of calls that satisfy the receiver-side constraint criteria, allowing the simulation to be interpreted in probabilistic rather than binary terms.

### 2.10 Estimating local density and neighbourhood statistics

The receiver-side constraint analysis requires an estimate of the local conspecific density *ρ*_*i*_(*t*) around each bat. Because the simulation tracks full three-dimensional positions, density is estimated using a fixed-radius neighbourhood count:

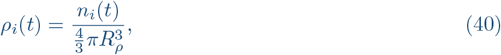

where *n*_*i*_(*t*) is the number of conspecifics within radius *R*_*ρ*_ of bat *i* (excluding *i*), and *R*_*ρ*_ is a user-defined kernel radius. This estimator is stable for heterogeneous groups and yields density in interpretable units (bats m^−3^). For the *post hoc* receiver-side analysis, the same value of *R*_*ρ*_ is used across all simulation runs to enable consistent comparison of density-dependent effects across parameter regimes.

The mean neighbourhood call rate 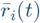 (Eq. (13)) is computed as the mean instantaneous call rate of conspecifics within the same density kernel. When only global statistics are required, 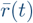 is taken as the mean call rate across all bats at time *t*.

### 2.11 Choice of vulnerable window and sensitivity checks

I define the vulnerable window as *W* = *τ* + *T*_*a*_ (Eq. (14)) because it captures two conservative conditions: interference concurrent with the outgoing call, and interference overlapping the earliest informative echo returns. In sensitivity analyses, I additionally tested narrower windows of the form

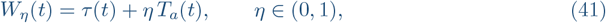

which represent cases where only a fraction of the echo delay is critical for behavioural updates (e.g. if early echo segments dominate distance estimation). Qualitative conclusions were robust across a broad range of *η*, with constraint boundaries shifting predictably as *η* decreased.

### 2.12 Acoustic parameterisation and simplifying assumptions

The overlap and dominance constraints require specifying acoustic parameters that vary across species and contexts. I therefore treat several terms as user-defined constants or scenario parameters.

#### Source level and frequency

In the feasibility model, *SL* (dB re 20 *µ*Pa at 1 m) and dominant frequency *f* determine attenuation through the absorption coefficient *α*(*f*) (Eq. (19)). When empirical values are unavailable for the simulated species, I report feasibility results across plausible ranges of (*SL, f*), rather than committing to a single species-specific calibration.

#### Directional gains

Directionality enters through *G*_tx_ and *G*_rx_ (Eq. (20)). In the main analysis, I present isotropic (worst-case) results by setting these gains to zero. Directional filtering can be included by restricting the effective solid angle Ω (Eq. (12)) and/or specifying gains as functions of angle, allowing the same framework to represent forward-directed emission beams and head/ear orientation effects.

#### Interferer statistics

I approximate interferer call onsets at the receiver as a Poisson process (Eq. (15)). This assumption is appropriate when many asynchronous emitters contribute and individual phase relations are not synchronised. I emphasise that this is a receiver-centric approximation: it does not require independence of call timing mechanisms across individuals, but assumes that the superposition of many call trains yields approximately Poisson arrival statistics at short time scales.

#### Stochastic geometry for dominance

The density-dependent dominance approximation (Eqs. (26)–(29)) treats conspecific positions as a spatial Poisson process within the sector volume. This provides a transparent first-order mapping between density and the expected proximity of the strongest (nearest) interferer. Where the simulated swarm exhibits structured spacing (e.g. lanes or local ordering), this approximation is conservative; in such cases, *r*_min_ may be replaced by the realised nearest-conspecific distance distribution extracted directly from simulation trajectories.

### 2.13 Outcome metrics and statistical summaries

I quantify swarm cohesion, safety, and sensory feasibility using the following outcome measures.

#### Cohesion and spacing

I report the distribution of nearest-neighbour distances *d*_*i*_(*t*), including mean, median, and selected quantiles over time and across individuals. Cohesion is summarised by (i) the time-resolved mean nearest-neighbour distance, and (ii) the dispersion of the swarm (e.g. mean distance to centroid), both computed from the full trajectory history.

#### Collision-risk rate

Collision risk is quantified as (i) the number of events where 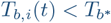 (Eq. (9)), and (ii) the number of TTC-violations (Eq. (11)), reported as absolute counts and normalised by total call events to obtain a rate per update.

#### Temporal-overlap fractions

I compute *P*_ov_(*t*) (Eq. (15)) for each bat at each call event using its local density estimate and neighbourhood call rate. I report the fraction of call events violating *P*_ov_ ≤ *p*_max_ and summarise how this fraction scales with *ρ* and with *k*_*r*_.

#### Level-dominance criterion

I compute whether the dominance criterion is satisfied (Eq. (29)) using scenario parameters (*SL, f, TS*, Δ, *q*, Ω, Θ_dir_). I report the fraction of call events for which neighbour echo dominance is expected to hold, and identify density ranges where dominance breaks down despite stable kinematics.

#### Joint constriant regimes

Finally, I quantify the proportion of call events satisfying both overlap and dominance constraints simultaneously (Eqs. (30)–(31)). This provides a direct estimate of the time spent in a regime where neighbour-based closed-loop updates are predicted to be reliable.

### 2.14 Variables and parameters

For clarity, model variables and parameters are summarised in two tables according to their functional role. SI Table 3 lists variables and parameters governing the swarm simulation and the closed-loop sonar control underlying individual flight behaviour, including kinematic state variables, responsivity-based timing parameters, and collision-risk metrics. SI Table 4 summarises variables and parameters introduced in the receiver-side feasibility model, which formalises temporal overlap and level-dominance constraints arising from conspecific acoustic interference.

**Table 3:** Variables and parameters governing swarm simulation and closed-loop sonar control.

| Symbol | Units | Description |
| --- | --- | --- |
| **Swarm state and kinematics** |  |  |
| $N$ | – | Number of bats in the swarm |
| $\mathbf{p}_i(t)$ | m | Position of bat $i$ |
| $\mathbf{h}_i(t)$ | – | Heading unit vector |
| $v_i(t)$ | $\text{m s}^{-1}$ | Flight speed |
| $v_0$ | $\text{m s}^{-1}$ | Initial speed |
| $\sigma_{\text{init}}$ | m | Initial spatial spread (SD per axis) |
| **Closed-loop sonar timing** |  |  |
| $d_i(t)$ | m | Distance to tracked neighbour |
| $c$ | $\text{m s}^{-1}$ | Speed of sound |
| $T_a(t)$ | s | Echo delay ($2d/c$) |
| $T_b(t)$ | s | Response interval ($k_r T_a$) |
| $k_r$ | – | Responsivity coefficient |
| $\text{IPI}(t)$ | s | Interpulse interval |
| $r(t)$ | Hz | Call rate |
| $\tau(t)$ | s | Call duration |
| $\tau_{\min}, \tau_{\max}$ | s | Bounds on call duration |
| $\kappa$ | – | Duration contraction factor |
| **Flight control** |  |  |
| $\alpha$ | – | Heading inertia |
| $\beta$ | – | Alignment weight |
| $\gamma$ | – | Cohesion weight |
| $\delta$ | – | Global drive weight |
| **Collision-risk metrics** |  |  |
| $T_b^*$ | s | Reaction-time threshold |
| TTC | s | Time-to-collision estimate |

**Table 4:** Variables and parameters governing temporal overlap, level dominance, and information transfer.

| Symbol | Units | Description |
| --- | --- | --- |
| **Local density and interference** |  |  |
| $\rho$ | bats $\text{m}^{-3}$ | Local swarm density |
| $\Omega$ | sr | Effective interference solid angle |
| $R$ | m | Interference radius |
| $V$ | $\text{m}^3$ | Interference volume |
| $\Lambda$ | $\text{s}^{-1}$ | Interferer arrival rate |
| $W$ | s | Vulnerable window ($\tau + T_a$) |
| $P_{\text{ov}}$ | – | Overlap probability |
| $p_{\text{max}}$ | – | Tolerable overlap threshold |
| **Acoustic propagation** |  |  |
| $SL$ | dB | Source level |
| $f$ | Hz | Call centre frequency |
| $\alpha(f)$ | $\text{dB m}^{-1}$ | Atmospheric absorption |
| $TL(r)$ | dB | Transmission loss |
| $\Theta_{\text{dir}}$ | dB | Audibility threshold (direct calls) |
| $\Theta_{\text{echo}}$ | dB | Echo detection threshold |
| **Level dominance** |  |  |
| $TS$ | dB | Neighbour target strength |
| $RL_{\text{echo}}$ | dB | Echo received level |
| $RL_{\text{max}}$ | dB | Strongest interferer level |
| $\Delta$ | dB | Required salience margin |
| $q$ | – | Reliability quantile |
| **Information transfer** |  |  |
| $I_t(n)$ | s | Information latency over $n$ hops |
| $I_v$ | $\text{m s}^{-1}$ | Information velocity ($c/2k_r$) |
| $\delta_0$ | – | Initial perturbation magnitude |
| $\alpha_{\text{att}}$ | – | Attenuation coefficient |
| $i_{\text{max}}$ | – | Maximum propagation range |

Simulation-specific parameters (e.g. number of agents *N*, initial spatial spread *σ*_init_, nominal flight speed *v*_0_, heading update weights *α, β, γ*, and *δ*, the physiological response threshold 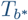, local density estimation radius *R*_*ρ*_, and bounds on individual speeds *v*_*i*_) are specified in the parameter grid and summary tables (Tables 1 and 2). Acoustic scenario parameters used to evaluate feasibility (including source level *SL*, dominant frequency *f*, atmospheric absorption *α*(*f*), effective interference geometry Ω, neighbour target strength *TS*, audibility and detection thresholds Θ_dir_ and Θ_echo_, tolerable overlap probability *p*_max_, reliability quantile *q*, and required salience margin Δ) are either fixed per analysis or systematically varied as indicated in the corresponding figure captions and supplementary materials.

## 3 RESULTS

To evaluate whether local echo-driven control alone can sustain swarm cohesion, I implemented a fully asynchronous simulation of echolocating bat aggregations. Each individual, modelled as an autonomous agent, independently emitted sonar calls based on its own sensory feedback loop, governed by the temporal precision theory. Unlike synchronous or discretely-timed models, this asynchronous design captures the natural variability in call emission and echo reception observed in real bats. Each bat adapts its call interval and duration according to its distance to the nearest neighbour ahead, while also updating its heading and speed based on local alignment and centre-of-mass cues. This framework closely mimics the real-world scenario of dense emergence swarms, where thousands of bats operate without central coordination yet maintain fluid, coherent group flight. The model allowed me to probe the emergent dynamics of such decentralised systems under systematically varied responsivity parameters (*k*_*r*_) and forward drive (*v*_0_), capturing how sonar-based control translates into collective stability, communication load, and collision risk.

I tested the impact of sonar responsivity and initial velocity on swarm dynamics. I conducted a systematic grid-based simulation sweep varying the responsivity coefficient (*k*_*r*_) and initial velocity (*v*_0_). Each simulation involved 50 bats flying for 60 seconds under closed-loop biosonar control. Output metrics included call rate, call duration, inter-bat distance, velocity, and collision risk events.

Despite the absence of any global coordination rule or prescribed spacing behaviour, all simulations produced globally cohesive swarm formations. As seen in Fig. 1, bats maintained forward progression with bounded lateral dispersion and low vertical spread, suggesting that local echo-based timing suffices to preserve group integrity over time.

**Figure 1:**
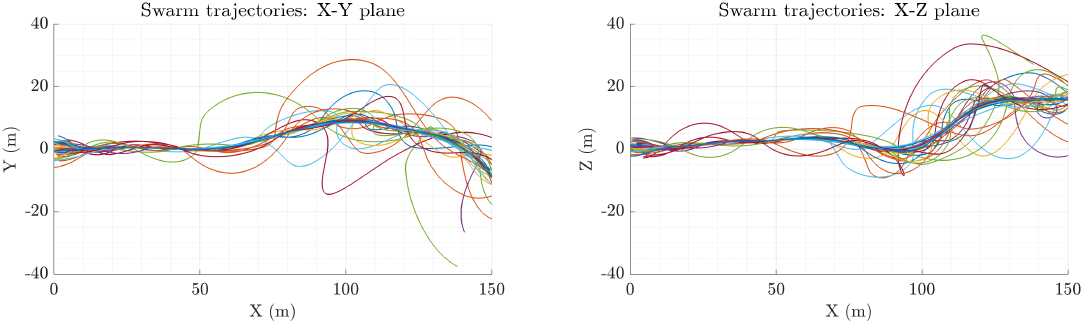
Swarm simulation showing the evolution of bat trajectories in the X–Y plane (left) and X–Z plane (right). Each line traces the path of one individual bat over the course of the simulation. The limited spread along the Z-axis demonstrates vertical cohesion, while horizontal positions (X–Y) exhibit local dispersion within the swarm structure. The forward progression along the X-axis reflects the effect of the global drive **g** applied to all bats. These patterns illustrate the stable, cohesive flow maintained by the simulated swarm under closed-loop sonar guidance.

### 3.1 Cohesive trajectories emerge from local echo-timed control

Across all simulated conditions, the swarm maintained sustained forward progression with bounded lateral dispersion and low vertical spread, despite the absence of any explicit spacing rule or global coordination. Representative trajectories illustrate stable collective flow in both the horizontal (X–Y) and vertical (X–Z) planes (Fig. 1), consistent with local echo-timed updates providing sufficient stabilising feedback at the level of individual interaction.

### 3.2 Call timing and spacing exhibit characteristic distributions

A representative run exhibited the predicted joint structure of call timing and spacing (Fig. 2). Call durations were frequently compressed towards the imposed physiological minimum (0.5 ms), consistent with frequent short-range interactions in which echo delays constrain available timing margins. Call rates spanned a wide dynamic range, reflecting continuous adjustment of IPI to neighbour distance under the closed-loop control law. Velocities remained tightly centred around the nominal forward speed, indicating that the asynchronous update scheme did not introduce drift or instability in locomotor output. Inter-individual distances exhibited a longtailed distribution with a modal spacing in the low-metre range, reflecting self-organised structure in which individuals intermittently converge and re-separate while preserving overall cohesion.

**Figure 2:**
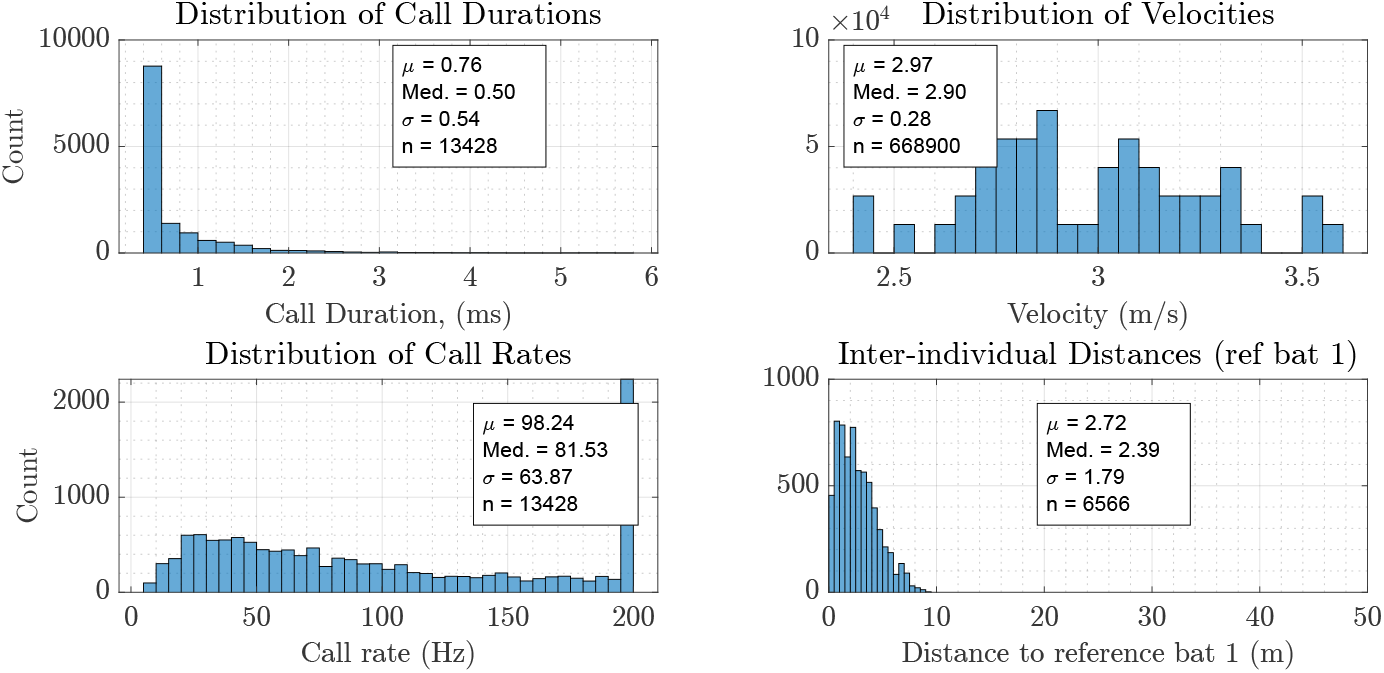
Summary statistics of a swarm simulation under the closed-loop echolocation behaviour (*N*_*bats*_ = 50, *T* = 5 s, *σ*_*init*_ = 2). **Top-left:** Distribution of call durations across all bats and time points. Most calls are dynamically compressed toward the physiological minimum (0.5 ms), reflecting close spacing within the swarm. **Top-right:** Distribution of flight velocities, showing stable speeds tightly centred around the initial target velocity (*µ* = 2.96 m/s). **Bottom-left:** Distribution of call rates, which adapt over a wide range (peak near 90 Hz) according to local inter-individual spacing. **Bottom-right:** Distribution of distances to a reference bat (bat 1), demonstrating self-organised swarm structure with a median spacing of ∼ 2.4 m and a long-tailed distribution from occasional forward or backward drift. These results confirm that closed-loop echolocation control can support stable, cohesive swarm flight in the absence of explicit central coordination.

### 3.3 Swarm-level parameter trends across *k*_*r*_ **and** *v*_0_

Swarm-level averages varied systematically with *k*_*r*_ and *v*_0_ (Fig. 3). Increasing *k*_*r*_ increased the effective response margin per unit echo delay and therefore lengthened IPI, yielding lower call rates and (via the call-duration contraction rule) longer typical call durations. Mean inter-individual spacing increased with larger *k*_*r*_, consistent with slower feedback promoting larger stable separations. Variation in *v*_0_ modulated these trends by shifting the kinematic context in which echo delays are sampled: faster forward motion produced larger typical separations in some regimes, particularly when paired with higher *k*_*r*_, indicating an interaction between locomotor pace and the timing budget available for closed-loop updates.

**Figure 3:**
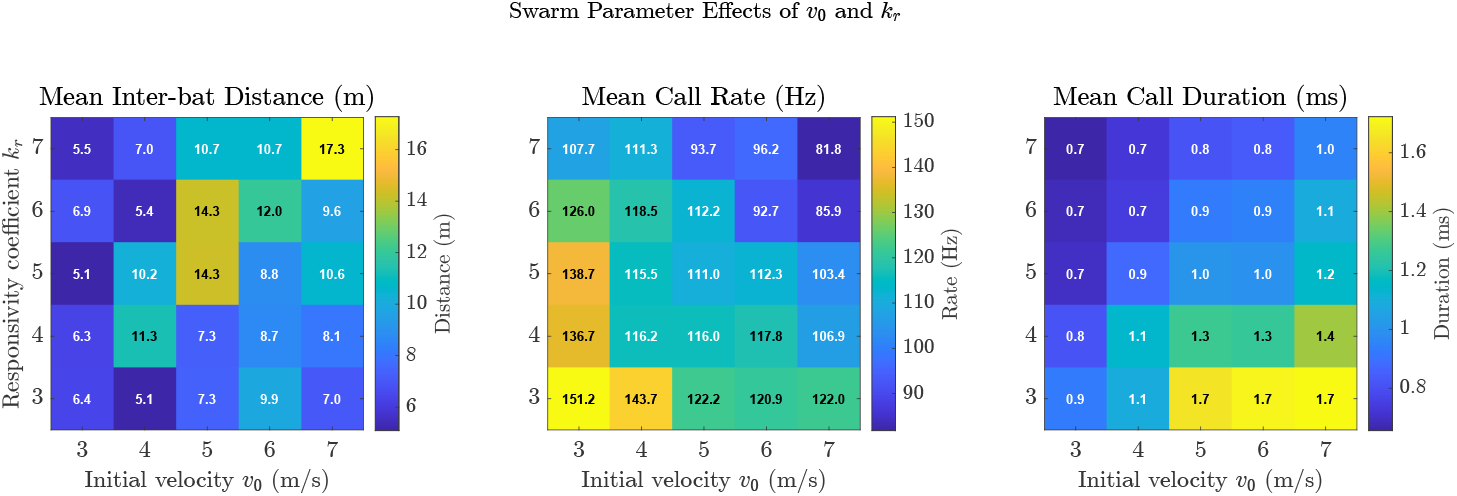
Swarm-level effects of initial velocity *v*_0_ and responsivity coefficient *k*_*r*_. Left: Mean inter-bat distance increases with higher *k*_*r*_ and peaks at intermediate–high *v*_0_ values, suggesting greater spacing emerges from slower responsiveness. Centre: Mean call rate declines with increasing *v*_0_ and *k*_*r*_, consistent with longer inter-pulse intervals at high flight speeds and echo delays. Right: Mean call duration shows a reciprocal trend to call rate, decreasing with slower responsiveness and faster speeds. Together, these trends highlight the coupled dynamics of biosonar timing and swarm spatial organisation.

Descriptive statistics for each simulation run are reported in Table 1, including summary measures of call rate, call duration, velocity, and neighbour distance across the full parameter grid.

### 3.4 Collision-risk regimes and safety margins

Collision-risk events were rare overall but non-zero in specific parameter regimes (Fig. 4). The temporal raster of warnings shows that risks occur intermittently rather than continuously, and are concentrated in combinations of higher forward speed and conservative responsivity that reduce the effective responsiveness of the loop relative to closing dynamics. The heatmap of normalised event rates reveals a distinct peak in collision-risk incidence at high *v*_0_ combined with high *k*_*r*_, whereas most regimes remain at very low rates. Pairwise risk margins further show that the majority of interactions retain positive margins (response time exceeding time-to-collision), with a small number of near-zero or negative margins marking boundary events where the closed-loop timing budget becomes insufficient.

**Figure 4:**
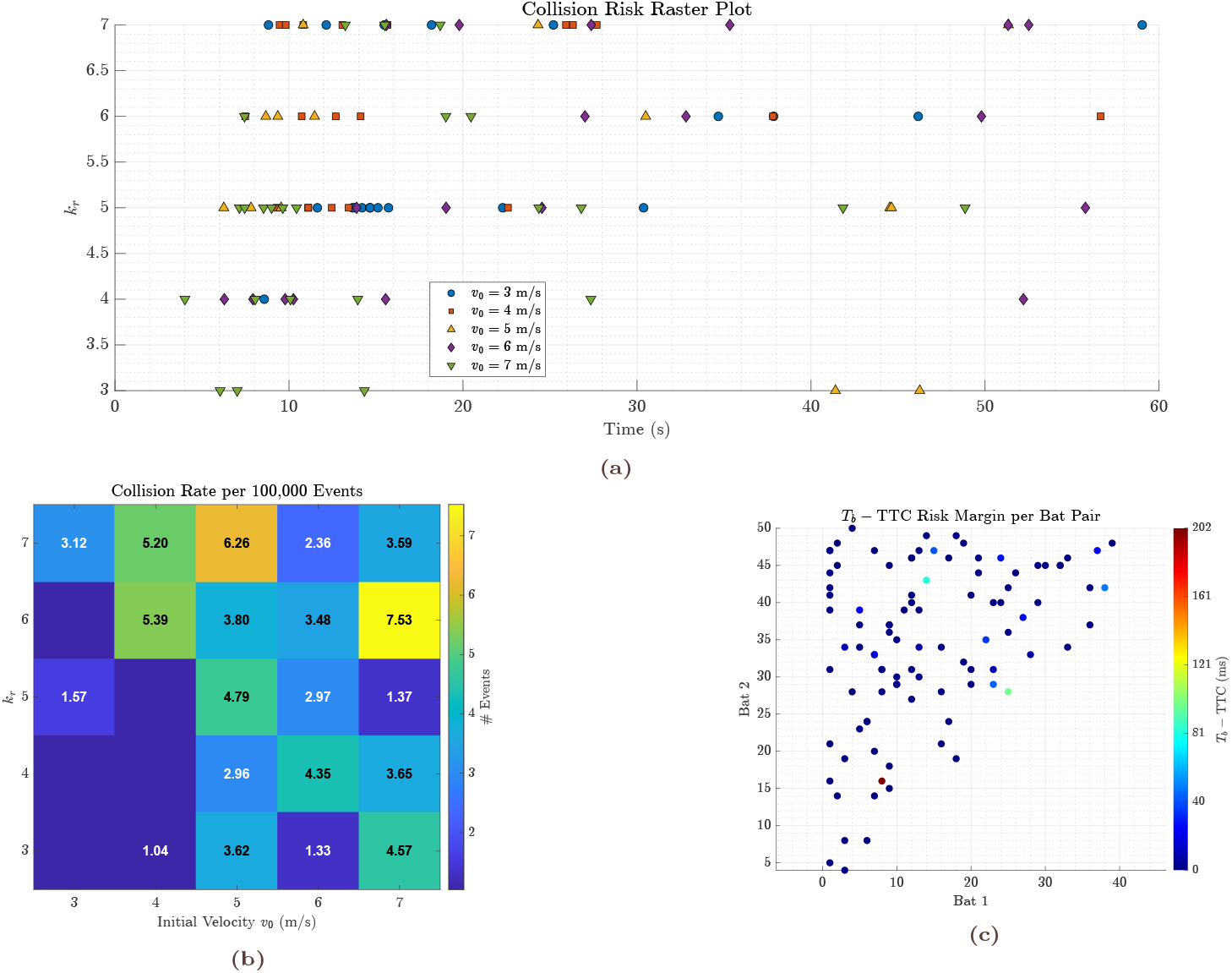
Comprehensive visualisation of collision risk events in the asynchronous swarm simulation with 50 bats and for 60 s. **Top:** Raster plot showing the occurrence of collision risk events over simulation time, stratified by responsivity coefficient (*k*_*r*_, vertical axis) and colour-coded by initial flight velocity (*v*_0_). Each point represents a collision warning flagged between a unique pair of bats. **Bottom left:** Heatmap showing the mean collision event rate (normalised per 100,000 update events) for each parameter combination of *k*_*r*_ and *v*_0_. Brighter values indicate higher collision frequencies. **Bottom right:** Pairwise risk margin plot across all bat pairs within a selected run, colour-coded by the difference between the response time (*T*_*b*_) and the time-to-collision (TTC). Positive margins indicate that the bats have sufficient time to respond before collision, while negative or small margins imply elevated risk. These analyses demonstrate that both *k*_*r*_ and *v*_0_ significantly influence collision dynamics, with higher speeds and lower responsivity values associated with increased collision likelihood.

Absolute and normalised collision event counts across the parameter grid are summarised in Table 2, showing that even in the highest-risk regimes, event rates remain low relative to total update events.

### 3.5 Receiver-side constraint satisfaction follows a probabilistic temporal-overlap boundary

Across the full parameter grid of responsivity (*k*_*r*_) and initial velocity (*v*_0_), receiver-side outcomes varied continuously rather than exhibiting discrete regimes. The fraction of call events for which both temporal-overlap and level-dominance constraints were satisfied ranged from approximately 2–25% across simulation runs (Fig. 5b). Temporal-overlap violations were frequent across all conditions, whereas dominance violations showed greater heterogeneity, indicating that the two constraints contribute differently to the overall probability of reliable receiver-side updating.

**Figure 5:**
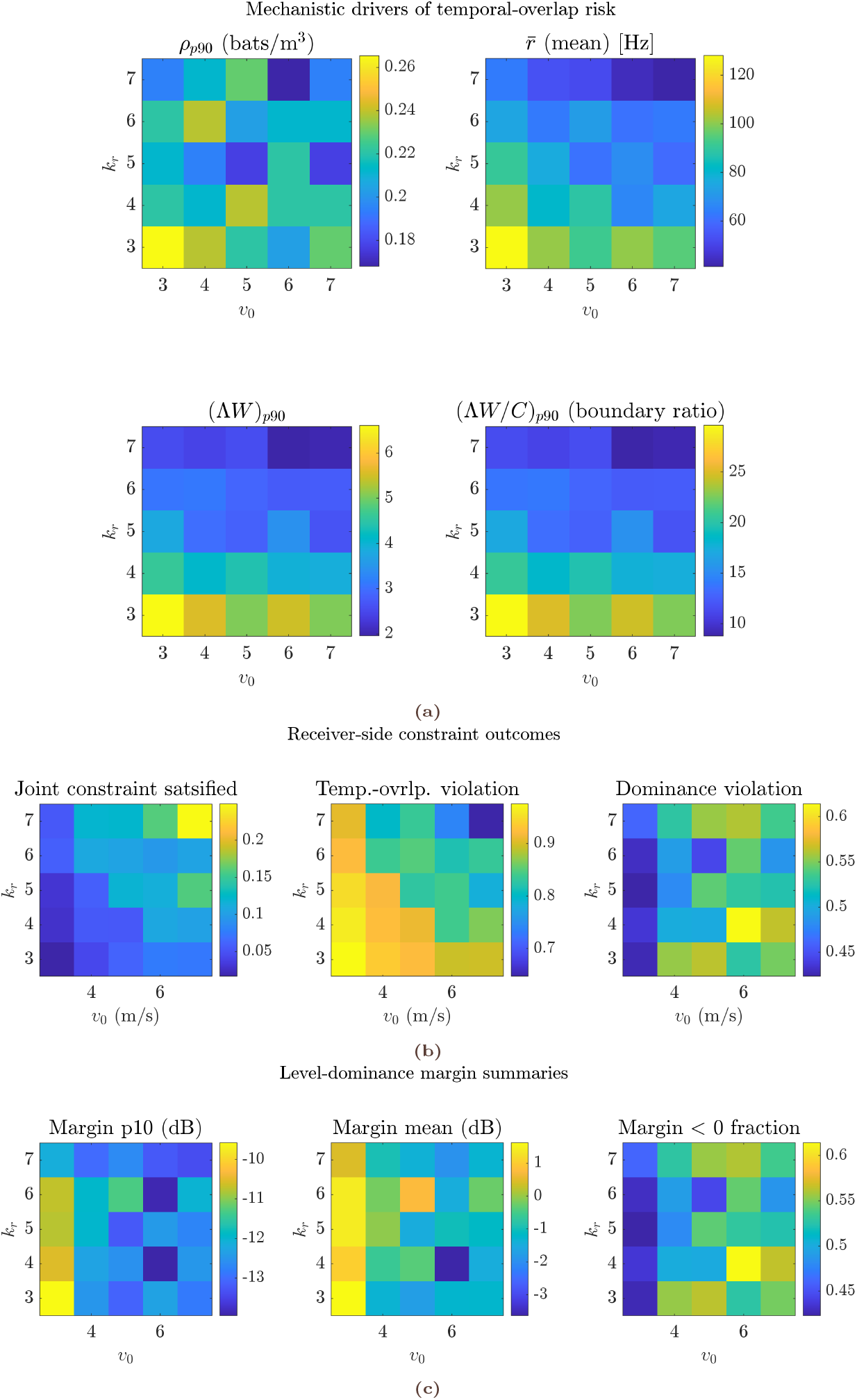
Receiver-side constraint structure and its mechanistic basis across the (*k*_*r*_, *v*_0_) grid. (a) Mechanistic maps decompose temporal-overlap risk into local density *ρ*_*p*90_, mean neighbourhood call rate 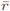, the compound information-load term (Λ*W*)_*p*90_, and the normalised boundary ratio (Λ*W/C*)_*p*90_ (with *C* = − ln(1 − *p*_max_)). These panels show that overlap risk is not set by density or call rate alone, but by their combined effect through Λ*W*. (b) Receiver-side outcomes summarise run-level fractions of joint constraint satisfaction, temporal-overlap violations, and dominance-criterion violations. Joint satisfaction varies smoothly across the grid and closely follows the boundary ratio rather than exhibiting sharp regime transitions. (c) Level-dominance margin summaries (10th percentile, mean, and fraction of margins *<* 0) reveal a broadly marginal masking regime, in which echo levels frequently approach or fall below competing call levels. Together, the panels demonstrate that receiver-side constraints in dense swarms operate probabilistically and continuously, shaped by compounded information load rather than by hard coordination or dominance thresholds.

Run-level boundary diagnostics revealed that temporal overlap is tightly organised by the compound information-load term (Λ*W*) evaluated at the 90th percentile of calls within each run (Fig. 6c). As (Λ*W*)_*p*90_ increased, the corresponding overlap probability *P*_ov,*p*90_ rose monotonically and approached unity, consistent with the analytical prediction in Eq. (37).

**Figure 6:**
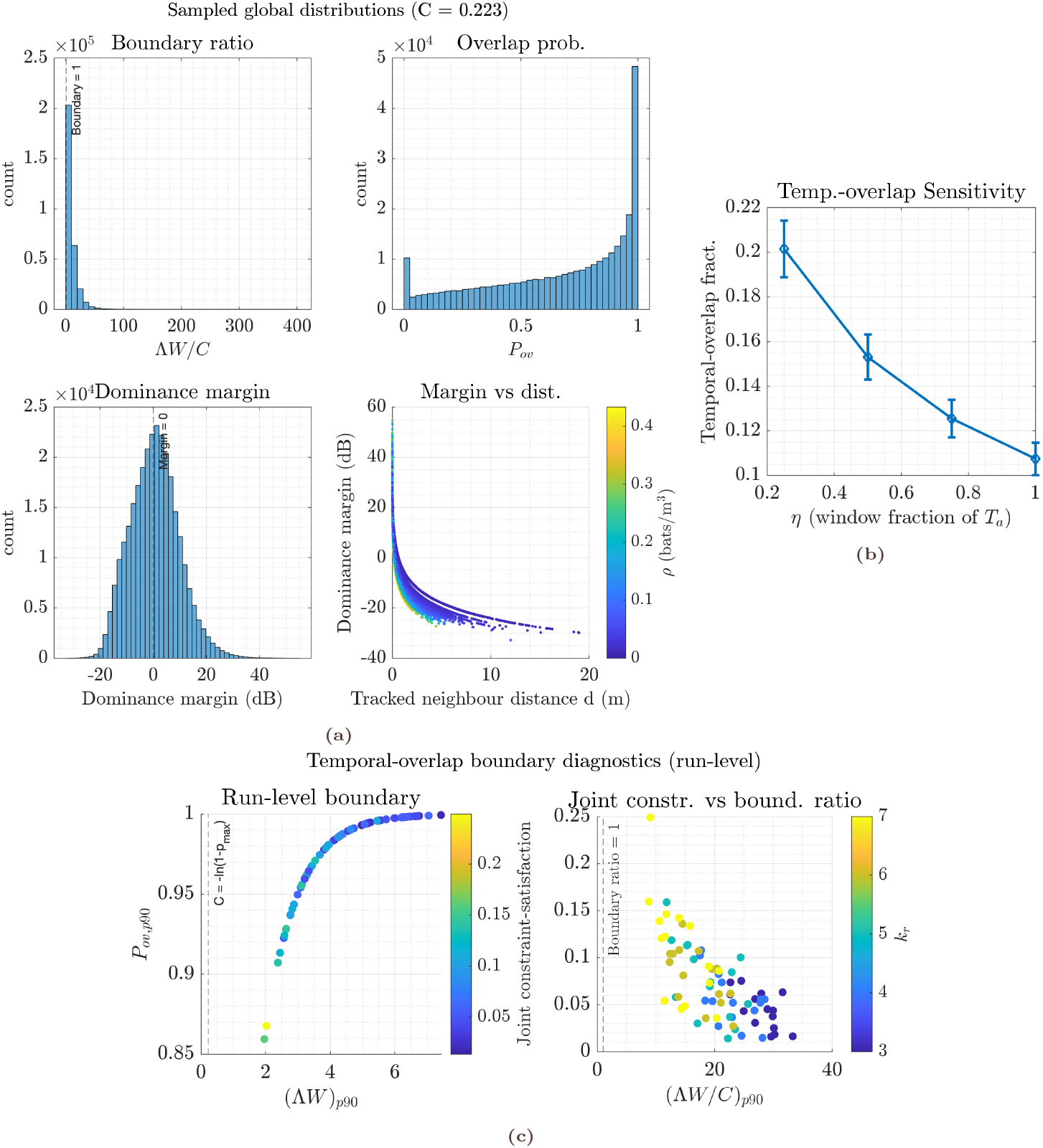
Temporal-overlap acts as a probabilistic receiver-side boundary and is robust to the temporal-window definition. (a) Global sampled distributions summarise the boundary ratio Λ*W/C*, the implied overlap probability *P*_ov_ = 1 − exp(− Λ*W*), the dominance margin, and the dependence of dominance margin on tracked-neighbour distance and local density *ρ*. These distributions show that most interactions operate near the overlap boundary and within a broadly marginal dominance regime. (b) Sensitivity of temporal-overlap constraint satisfaction to the temporal window definition. Increasing the window fraction *η* in *W* = *τ* + *ηT*_*a*_ produces a smooth, monotonic decrease in the fraction of call events satisfying the overlap criterion, without qualitative changes in ordering across runs. (c) Run-level boundary diagnostics demonstrate that *P*_ov,*p*90_ increases monotonically with (Λ*W*)_*p*90_ and that joint constraint-satisfaction declines systematically with the normalised boundary ratio (Λ*W/C*)_*p*90_ (dashed line indicates Λ*W/C* = 1). Together, these panels show that temporal overlap imposes a graded, probabilistic constraint on receiver-side processing rather than a sharp coordination threshold.

Normalising by the boundary constant *C* = − ln(1 − *p*_max_) collapsed runs onto a single diagnostic axis: the fraction of call events satisfying both receiver-side constraints declined systematically once the boundary ratio (Λ*W/C*)_*p*90_ exceeded unity (Fig. 6c). Importantly, this decline was gradual rather than abrupt, demonstrating that receiver-side constraints act probabilistically rather than imposing hard behavioural limits.

### 3.6 Mechanistic drivers of temporal-overlap constraint satisfaction

Decomposing the compound load term revealed how temporal-overlap constraint satisfaction emerges from interacting ecological and behavioural variables (Fig. 5a). High local densities (*ρ*_*p*90_) and elevated neighbourhood call rates 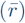 both increased temporal overlap probability, but neither variable alone predicted run-level outcomes. Instead, receiver-side constraint satisfaction aligned most consistently with the combined term (Λ*W*)_*p*90_ and its normalised boundary ratio (Λ*W/C*)_*p*90_.

Across the parameter grid, runs with comparable densities could differ substantially in overlap outcomes depending on call timing and responsivity, illustrating that temporal organisation modulates the effects of crowding. Conversely, even moderate densities produced high overlap probabilities when coupled with elevated call rates and extended effective temporal windows.

These patterns confirm that temporal overlap is governed by a compounded information-load constraint rather than by static spatial density alone.

### 3.7 Level dominance operates in a marginal probabilistic regime

Analysis of level dominance revealed that masking is pervasive rather than exceptional (Fig. 5c). The 10th-percentile dominance margin was negative across much of the parameter space, indicating that a substantial fraction of call events experienced interferer levels exceeding echo levels. Mean dominance margins clustered near 0 dB, and in many conditions more than half of calls fell below the dominance criterion.

Global distributions pooled across runs further illustrated this marginal regime (Fig. 6a). Dominance margins were approximately symmetric around zero with a long positive tail, implying that strong dominance events occur but are rare, whereas near-threshold or negative margins are common. Dominance margin declined steeply with increasing tracked-neighbour distance, particularly at higher local densities, highlighting the joint influence of spatial configuration and crowding on masking risk.

These results indicate that receiver-side processing cannot rely on consistent level dominance. Instead, behavioural updating must accommodate frequent masking events and operate within probabilistic tolerance limits rather than enforcing dominance as a strict requirement.

### 3.8 Robustness to temporal-window assumptions

Sensitivity analysis confirmed that the observed receiver-side constraint structure is robust to the definition of the effective temporal window defined in Eq. (41) (Fig. 6b). Increasing the window fraction *η* produced a smooth, monotonic decline in the fraction of call events satisfying the temporal-overlap constraint, as expected from the analytical form of *P*_ov_. No qualitative changes in run ordering were observed, and the probabilistic boundary relationship remained intact.

This robustness demonstrates that the framework does not depend on finely tuned parameter choices, but instead reflects a general constraint imposed by call timing, interaction geometry, and local information load.

## 4 DISCUSSION

Cohesive animal groups must continuously update spacing and headings while operating under finite sensing delays, processing time, and motor latency. In echolocating bats, these constraints are unusually tight because sensory acquisition is *self-generated* and inherently closed-loop: call emission, echo return, processing, and motor update must remain temporally compatible if neighbour-based interactions are to remain stable. Building on an asynchronous swarm simulation with echo-timed updates, the present work adds an explicit receiver-side feasibility layer that turns the social “cocktail-party” problem into two analytically tractable constraints: (i) *temporal overlap* within a biologically vulnerable window, and (ii) *level dominance* of the tracked neighbour echo over the strongest competing conspecific signal. Together, these constraints move the question from “can echoes be detected under masking?” to “when is a neighbour-tracking loop dynamically self-consistent as density increases?”

### 4.1 Closed-loop timing links neighbour geometry to acoustic adaptation

The asynchronous implementation is important because it avoids imposing global synchrony: each bat updates only when its own sensory loop completes, with inter-pulse interval (IPI) and call duration emerging from local echo delay and the responsivity rule. This captures a central biological reality of dense emergence streams: even if many individuals occupy the same volume, their sensing and actuation are not clocked to a shared schedule. In this setting, cohesive motion can arise from a minimal interaction rule (nearest forward neighbour plus weak alignment/cohesion drives) *provided that* the loop remains temporally feasible. The model therefore frames cohesion as an emergent property of many locally stable temporal loops rather than as an outcome of explicit spacing objectives or global information.

A robust qualitative outcome of the closed-loop rule is the tight coupling between echo delay, call duration, and call rate: as neighbour distance decreases, echo delay shortens, calls contract towards a physiological minimum, and call rate increases. This pattern aligns with the empirical signature of crowded flight in multiple systems, including the onboard emergence results of [15] and laboratory work on cluttered tracking in [27]. Field observations of swarm-size-dependent emission rate increases [28] and group-flight call shortening [29, 30] are consistent with the same principle: local geometry (and thus echo delay) is sufficient to drive broad acoustic reconfiguration without invoking explicit “anti-jamming” strategies. Socially modulated beam/timing adjustments reported in paired-flight contexts [31] further support the idea that bats actively tune what counts as the most relevant neighbour in the loop, not merely the raw detectability of any echo.

### 4.2 Probabilistic receiver-side feasibility and its behavioural consequences

The new contribution of the feasibility framework is to show that call adjustments are simultaneously *solutions* to, and *drivers* of, interference. Two effects are central.

#### Temporal-overlap constraint as a density–timing boundary

When conspecific onsets arrive at the receiver with aggregate rate 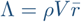 (local density *ρ*, effective interference volume *V*, mean call rate 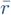), the probability that at least one onset intrudes into the vulnerable window *W* = *τ* + *T*_*a*_ is

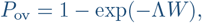

yielding the feasibility requirement *P*_ov_ ≤ *p*_max_, or equivalently,

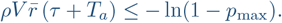

This makes a concrete prediction: for fixed geometry (hence *V*) and target distance (hence *T*_*a*_), increasing call duration *τ* or call rate 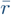 pushes the system towards an overlap-dominated regime even if echo levels remain high. Importantly, overlap is not intrinsically disruptive; it becomes limiting when the *timing demands of dominant interactions* exceed what can be processed within stable update windows.

#### Level dominance as a probabilistic constraint on which neighbours can drive behaviour

In dense scenes, behaviour is unlikely to be controlled by the total number of conspecifics present. Instead, a small subset of *level-dominant* individuals disproportionately shapes the received sound field. This motivates the dominance requirement

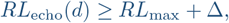

where *RL*_echo_(*d*) is the neighbour echo at distance *d, RL*_max_ is the strongest competing direct call among relevant interferers, and Δ is a salience margin. Expressing *RL*_max_ in terms of the nearest (or strongest) interferer distance distribution under a sector-limited spatial point process yields an explicit mapping from density to the expected dominance failure rate. The consequence is a second sharp prediction: beyond a density-dependent boundary, neighbour tracking is not limited by “hearing in noise” in general, but by the inability of *the tracked echo* to remain reliably more informative than the strongest competitor.

#### Trade-offs and why empirical call changes need not reduce interference

The frame-work provides a mechanistic interpretation for why bats in strong interference often call louder, longer, and more frequently [12] without necessarily achieving reduced jamming: increasing *SL* can improve *RL*_echo_ and help satisfy dominance, but it also increases the interference radius (hence *V*) and therefore increases Λ and overlap probability; increasing *τ* increases echo energy but expands *W*; increasing call rate increases update rate (reducing state staleness) but increases Λ by construction. These opposing effects formalise why call adjustments alone cannot guarantee stable neighbour-based cohesion at very high densities, and why additional degrees of freedom (directionality, spatial filtering, neighbour selection rules, or cue substitution) may become necessary.

The receiver-side analyses provide a mechanistic account of how interaction density constrains closed-loop neighbour tracking in echolocating swarms. Rather than imposing binary conditions for perceptual success or failure, temporal overlap and level dominance define *probabilistic limits* on how frequently usable sensory updates are available to support behavioural control. Across the (*k*_*r*_, *v*_0_) parameter space, neither constraint produced sharp regime transitions. Instead, the fraction of call events capable of supporting reliable neighbour-based tracking declined smoothly with increasing interaction load.

Temporal overlap was organised by the compound load term Λ*W*, integrating local density, neighbourhood call rate, call duration, and echo delay. Overlap probability followed the analytical form *P*_ov_ = 1 − exp(−Λ*W*), and run-level diagnostics showed a systematic decline in joint constraint satisfaction once (Λ*W/C*)_*p*90_ approached unity. Importantly, this transition was gradual rather than abrupt, indicating that overlap limits behavioural updating statistically rather than enforcing a hard coordination threshold. Overlapping calls therefore do not directly imply perceptual failure; instead, they reduce the availability of timely feedback needed to sustain closed-loop control.

Level dominance exhibited a complementary structure. Dominance margins clustered near zero, with a substantial fraction of call events experiencing negative margins in which direct conspecific calls exceeded echo levels. These marginal conditions were common even in regimes where flight kinematics and call timing remained stable, indicating that neighbour tracking in dense groups does not rely on persistent echo dominance. Instead, reliable control must tolerate frequent masking and operate through redundancy across calls and interactions.

Together, these constraints define a receiver-side landscape in which cohesion is maintained as long as a sufficient density of temporally and acoustically viable updates is available. Behavioural control degrades when interaction load reduces this fraction below a critical level, even though individual echoes may remain detectable. Breakdown therefore reflects an exhaustion of response feasibility rather than a loss of sensory access.

This probabilistic structure has direct behavioural implications. A focal bat need not, and likely cannot, continuously track a single nearest neighbour. Experimental data of interpulse interval dynamics show alternating IPI swings, consistent with sequential attention to different objects of relevance rather than persistent fixation [16, 32, 33]. In dense swarms, where relative velocities and interaction partners change unpredictably, such alternating tracking becomes essential—an expectation in-line with previous studies [34–36]. When a particular echo is rendered uninterpretable by overlap or insufficient dominance, subsequent echoes may still carry sufficient information to support the next behavioural update. Closed-loop stability therefore depends on the rate at which informative updates occur, not on uninterrupted dominance of any single echo.

Within this framework, the joint constraint-satisfaction probability acts as the effective control variable. It governs how densely packed neighbours can be while still permitting timely responses, and it links individual sensory processing limits to collective behaviour. Species-specific differences in responsivity and time budget determine how many peers can be accommodated within this probabilistic window.

Crucially, this perspective reframes the role of acoustic interference in swarms. Overlap and masking are unavoidable in dense groups, but they do not necessitate invoking jamming as a failure mode. Instead, interference shapes the statistics of information availability, and behavioural responses to crowding emerge from attempts to maximise the probability of receiving relevant echo information within a limited time window. Because each bat experiences a unique local feasibility landscape, such adjustments introduce perturbations that propagate through the group via changes in relative position and interaction timing. The same closed-loop principles therefore explain both stable cohesion at high update rates and the emergence of spreading, reorganisation, or fragmentation when probabilistic feasibility becomes locally exhausted.

Thus, temporal overlap and dominance do not constrain perception directly; they regulate the flow of actionable information that sustains collective stability.

### 4.3 Relation to masking, jamming, and swarm sensing models

The “sonar cocktail party nightmare” model of Beleyur & Goerlitz (2019) [13] explicitly simulates many biologically relevant contributors to masking and derives predictions about when neighbour echoes should remain detectable in groups. Their deliberate choice to hold call parameters fixed and apply simplified reception criteria makes the conclusions conservative and clean for perceptual feasibility. The present framework complements that work by addressing a different question: whether a *closed-loop neighbour-tracking rule* can remain self-consistent under increasing density. Temporal overlap and dominance convert masking from an outcome into feasibility inequalities, thereby predicting *density-dependent breakdown* (or required co-adjustments of *τ*, IPI, and *SL*) even in parameter regions where “some detection” might still be possible in an open-loop sense.

#### 4.3.1 Empirical adjustments as constraints to be explained

Amichai et. al. (2015) [12] show that bats under severe conspecific-like interference primarily increase intensity, duration, and rate, while spectral shifts are modest and not reliably deconflicting. Within the present framework, these responses emerge as partial attempts to recover echo salience (dominance), but they can simultaneously exacerbate temporal overlap by inflating *W* and Λ through increased *V* and 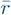. The key interpretive shift is that “louder and longer” is not a universal remedy; it is a strategy with a predictable failure mode when density pushes the system into an overlap-dominated regime.

#### 4.3.2 Detailed sensorimotor simulation of jamming versus analytical feasibility limits

Mazar & Yovel (2020) [14] evaluated whether a spectral jamming avoidance response improves reception using a rich agent-based sensorimotor model and multiple receiver criteria. Their negative result (JAR is not reliably beneficial) is naturally compatible with the feasibility view: if the dominant failure mode is intrusion into a short vulnerable window (i.e. Λ*W* becomes large), then frequency manoeuvres cannot prevent disruption of the timing backbone. Likewise, dominance depends on spatial statistics and attenuation, not only on spectral placement. The two approaches therefore address different explanatory layers: Mazar & Yovel (2020) [14] quantify outcome frequency given assumed strategies; the present framework predicts which regions of strategy space are feasible at all as density increases.

#### 4.3.3 Beyond self-echo attribution: soundscape navigation as an alternative regime

The soundscape-gradient hypothesis of Vanderelst & Peremans (2025) [18] proposes that obstacle avoidance and collision prevention can occur without isolating self-generated echoes, by exploiting collective amplitude gradients. This is not in conflict with loop-feasibility constraints. Instead, it provides a plausible regime *beyond* the feasibility boundary: when overlap and/or dominance cannot be jointly satisfied, strategies that do not require self-echo attribution become selectively advantageous. The present framework contributes a principled criterion for when such a regime shift should occur, expressed in measurable ecological variables (local density, call rate statistics, attenuation-limited interaction geometry, and required salience margin). The empirical evidence against this prediction remains to be established.

### 4.4 Empirical anchors: feasibility regimes and collision risk as timing failures

Goldshtein et al. (2025) [15] provide rare field access to the emergence problem by combining reconstructed trajectories of many individuals with onboard microphones, alongside a sensorimotor model linking interference to movement. Two aspects are particularly informative for the feasibility perspective.

First, their key empirical trajectory—high masking near the roost followed by a rapid drop as the emergence stream spreads—is consistent with the interpretation that spatial spreading restores feasibility by reducing *ρ* and thus Λ and *RL*_max_. Second, their results suggest an implicit separation of functions: even when a large fraction of potential echoes are jammed, *collision risk can remain low* because near-field interactions benefit from steep distance dependence of echo attenuation (∝ *d*^−4^) and because redundancy across calls allows occasional unjammed measurements to suffice. In the dominance framework, this corresponds to a regime where *RL*_echo_(*d*) can still exceed *RL*_max_ for very small *d* even when cohesion-maintaining tracking at larger interaction distances is already compromised. The feasibility model therefore refines the interpretation: at extreme densities, bats may retain reliable *local avoidance* while losing reliable *broader neighbour-based cohesion*, motivating either continued spreading, stricter neighbour selection, increased directionality, or a transition to soundscape-based cues.

In the simulation, collision risk emerges when the time-to-collision (TTC) becomes shorter than the effective behavioural response interval. This places collision avoidance within the same timing logic as cohesion: safety is maintained when updates arrive fast enough and remain interpretable. The feasibility framework adds a critical nuance: even if kinematics would permit avoidance, overlap and dominance can prevent the loop from delivering a reliable update at the required moment. This predicts a characteristic failure pattern at high density: collision risk need not increase smoothly with crowding; it can remain low across a wide range and then rise sharply once the system crosses an overlap- or dominance-driven feasibility boundary. This is consistent with the general claim that fragmentation or instability can appear without any change in “motivation” or control objectives, simply because the loop cannot be closed in time.

### 4.5 Information transfer and interaction dominance: what propagates through the swarm?

The information-transfer model in the present study shows that perturbations propagate through chains of neighbour interactions with an effective velocity governed primarily by the responsivity parameter (and sound speed), while attenuation limits the spatial extent over which any disturbance remains behaviourally relevant. The dominance concept tightens the biological interpretation: what propagates is not “information from all neighbours”, but updates driven by whichever interactions are *currently dominant* in level and timing. As density rises, dominance becomes more volatile (the identity of the strongest competitor changes faster), which can shorten the effective communication radius even if the kinematic coupling remains intact. This provides a route by which dense groups can remain cohesive yet become locally “myopic”, relying on a smaller subset of near-field interactions.

### 4.6 Implications for biomimetic swarm control

For engineered swarms, the present framework motivates a design principle distinct from optimisation-based bat-inspired heuristics [37]: stable cohesion can be achieved with decentralised, echo-timed control *only if* the sensing–actuation loop remains feasible under interference. The overlap and dominance inequalities therefore function as explicit engineering constraints that can be monitored online (via estimates of local density, event rates, and received-level statistics) to trigger regime shifts: e.g. from neighbour-echo tracking to coarser neighbour selection, stronger directionality (reducing Ω and *V*), or global cues. This aligns with broader arguments for biologically grounded sensory control in aerial autonomy [38–40], while adding a quantitative mechanism for when and why decentralised strategies should fail.

### 4.7 Functional implications: probabilistic sensing, anticipation, and swarm cohesion

The receiver-side results developed here reframe swarm cohesion as an emergent consequence of probabilistic sensory feasibility rather than interference avoidance or explicit coordination. Temporal overlap and level dominance do not define binary limits on perception; instead, they regulate the *rate* at which actionable sensory updates become available to sustain closed-loop control. Cohesion is maintained as long as this update rate remains sufficient, even though many individual call–echo events are masked.

This structure explains how dense echolocating swarms remain stable despite pervasive acoustic interference. Rather than requiring uninterrupted access to clean, dominant echoes, bats operate in a regime where informative updates are noisy. Behavioural control is therefore supported by statistical sufficiency across time, not by continuous perceptual clarity. This naturally implies alternating attention among nearby conspecifics, consistent with observed interpulse-interval dynamics, and allows neighbour relevance to be transient rather than fixed.

Notably, the results also admit an anticipatory complement to sensory processing [41]. Because relative velocities within emerging swarms are typically low, echo arrival times evolve predictably across successive calls. Even when individual echoes are partially masked or temporally overlapped, expectations based on prior echo timing may help constrain interpretation, reducing the effective impact of interference, resembling *entrainment* [42]. Such anticipation does not eliminate overlap or masking, but it further increases the probability that usable information can be extracted from a noisy stream.

Within this framework, acoustic interference is not a failure mode but a background condition that shapes the statistics of information availability. Local behavioural adjustments that improve echo timing or dominance increase the probability of timely updates and therefore become selectively favoured. Because each bat experiences a distinct local feasibility landscape, these adjustments propagate through the swarm, producing spreading, reorganisation, or fragmentation when probabilistic limits are locally exceeded.

Overall, the findings support a view of swarm cohesion as a sensory-driven, self-regulating process governed by probabilistic sensory constraints on information flow.

## SUMMARY OF REVISIONS

The revised manuscript with the updated title incorporates substantial conceptual, methodological, and interpretive changes in response to reviewer feedback. The major revisions are summarised below.

### 1. Probabilistic receiver-side constraints

The central conceptual update is the inclusion and formulation of temporal overlap and level dominance as *probabilistic feasibility boundaries* rather than hard perceptual failure modes. The revised analysis demonstrates that receiver-side constraints regulate the *frequency* of usable sensory updates, aligning swarm cohesion with graded information availability rather than strict coordination or interference avoidance.

### 2. Introduction of a dimensionless temporal-overlap boundary ratio

The temporal overlap model has been extended by introducing a normalised boundary ratio, Ξ = Λ*W/C*, where *C* = − ln(1 − *p*_max_). This reformulation clarifies the interpretation of overlap risk, enables direct comparison across parameter regimes, and reveals a common probabilistic boundary organising receiver-side feasibility across simulations.

### 3. Mechanistic decomposition of interaction load

New mechanistic analyses explicitly decompose overlap risk into local density, neighbourhood call rate, and timing structure, demonstrating that no single variable predicts feasibility in isolation. This addresses concerns regarding density-only or rate-only explanations and shows that compounded information load governs receiver-side limits.

### 4. Explicit treatment of marginal masking regimes

The revised manuscript adds dominance-margin analyses showing that most interactions operate near 0 dB dominance, with frequent masking events even in stable behavioural regimes. This reframes acoustic interference as a persistent background condition rather than a failure state and avoids reliance on jamming-based interpretations.

### 5. Behavioural interpretation via diffuse attention and closed-loop stability

The discussion has been substantially expanded to link probabilistic feasibility to behavioural strategies, including alternative neighbour tracking and redundancy across calls. This interpretation integrates the receiver-side results with the broader closed-loop framework of the paper and explains how cohesion can be maintained despite frequent overlap and masking.

Together, these revisions shift the manuscript toward a probabilistic, receiver-centric framework in which collective stability emerges from the statistics of sensory update availability rather than from group-level coordination or uninterrupted echo dominance.

## ACKNOWLEDGEMENTS

I gratefully acknowledge the support and insightful guidance of my doctoral supervisor Uwe Firzlaff, and thank my colleagues for their valuable discussions, encouragement, and feedback throughout the development of this work. I also thank the two anonymous reviewers for their constructive and thoughtful comments, which prompted substantial improvements to the scope, clarity, and ecological interpretation of the study.

## CODE AND DATA AVAILABILITY

The simulation code, experimental data, and analysis scripts are available in a repository archieved at DOI: https://doi.org/10.6084/m9.figshare.29482268.v1 Additionally, a GitHub repository is available with the code at: https://github.com/raviumadi/Swarming_Dynamics

## ETHICS STATEMENT

This study did not involve human participants or live animals. All analyses and simulations were conducted computationally, and no ethical approval was required.

## COMPTETING INTERESTS

The author declares no competing interests.

## A SUPPLEMENTARY INFORMATION

### A.1 Pseudocode

#### Algorithm 1

Asynchronous Bat Swarm Simulation

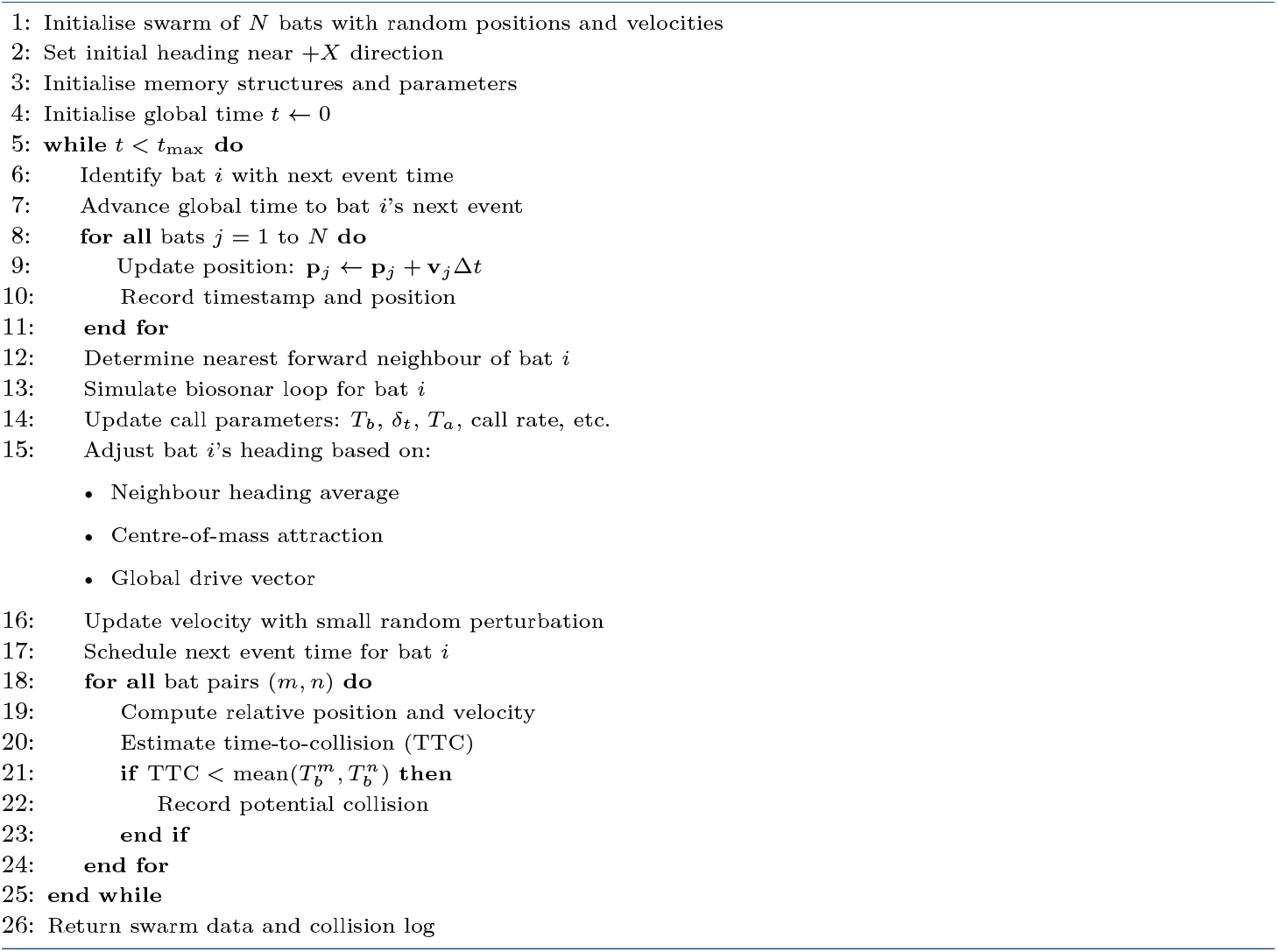

### A.2 Variable Lists

## REFERENCES

1. Couzin, I. D., Krause, J. E. N. S., James, R. I. C. H. A. R. D., Ruxton, G. D. & Franks, N. R. Collective Memory and Spatial Sorting in Animal Groups. Journal of Theoretical Biology 218, 1–11. ISSN: 0022-5193. doi:10.1006/jtbi.2002.3065 (Sept. 7, 2002).

2. Krause, J. & Ruxton, G. D. Living in Groups ISBN: 978-0-19-850817-5. doi:10.1093/oso/9780198508175.001.0001 (Oxford University Press, Oct. 17, 2002).

3. Vicsek, T., Czirók, A., Ben-Jacob, E., Cohen, I. & Shochet, O. Novel Type of Phase Transition in a System of Self-Driven Particles. Physical Review Letters 75, 1226–1229. doi:10.1103/PhysRevLett.75.1226 (Aug. 7, 1995).

4. Boonman, A., Fenton, B. & Yovel, Y. The Benefits of Insect-Swarm Hunting to Echolocating Bats, and Its Influence on the Evolution of Bat Echolocation Signals. PLOS Computational Biology 15, e1006873. ISSN: 1553-7358. doi:10.1371/journal.pcbi.1006873 (Dec. 12, 2019).

5. Tsoar, A. et al. Large-Scale Navigational Map in a Mammal. Proceedings of the National Academy of Sciences 108, E718–E724. doi:10.1073/pnas.1107365108 (Sept. 13, 2011).

6. Cvikel, N. et al. Bats Aggregate to Improve Prey Search but Might Be Impaired When Their Density Becomes Too High. Current biology: CB 25, 206–211. ISSN: 1879-0445. doi:10.1016/j.cub.2014.11.010. PMID: 25578909 (Jan. 19, 2015).

7. Warnecke, M., Chiu, C., Engelberg, J. & Moss, C. F. Active Listening in a Bat Cocktail Party: Adaptive Echolocation and Flight Behaviors of Big Brown Bats, Eptesicus Fuscus, Foraging in a Cluttered Acoustic Environment. Brain, Behavior and Evolution 86, 6–16. ISSN: 1421-9743. doi:10.1159/000437346. PMID: 26398707 (Sept. 2015).

8. Bee, M. A. & Micheyl, C. The “Cocktail Party Problem”: What Is It? How Can It Be Solved? And Why Should Animal Behaviorists Study It? Journal of comparative psychology (Washington, D.C. : 1983) 122, 235–251. ISSN: 0735-7036. doi:10.1037/0735-7036.122.3.235. PMID: 18729652 (Aug. 2008).

9. Griffin, D. R. Listening in the Dark: The Acoustic Orientation of Bats and Men xviii, 413. xviii, 413 (Yale Univer. Press, Oxford, England, 1958).

10. Schnitzler, H.-U. & Kalko, E. K. V. Echolocation by Insect-Eating Bats. BioScience 51, 557. ISSN: 0006-3568. doi: 10.1641/0006-3568(2001)051[0557:ebieb]2.0.co;2 (2001).

11. Brumm, H. & Slabbekoorn, H. in Advances in the Study of Behavior 151–209 (Academic Press, Jan. 1, 2005). doi:10.1016/S0065-3454(05)35004-2.

12. Amichai, E., Blumrosen, G. & Yovel, Y. Calling Louder and Longer: How Bats Use Biosonar under Severe Acoustic Interference from Other Bats. Proceedings. Biological Sciences 282, 20152064. ISSN: 1471-2954. doi:10.1098/rspb.2015.2064. PMID: 26702045 (Dec. 22, 2015).

13. Beleyur, T. & Goerlitz, H. R. Modeling Active Sensing Reveals Echo Detection Even in Large Groups of Bats. Proceedings of the National Academy of Sciences of the United States of America 116, 26662–26668. ISSN: 1091-6490. doi:10.1073/pnas.1821722116. PMID: 31822613 (Dec. 26, 2019).

14. Mazar, O. & Yovel, Y. A Sensorimotor Model Shows Why a Spectral Jamming Avoidance Response Does Not Help Bats Deal with Jamming. eLife 9 (eds King, A. J., Vanderelst, D. & A Simmons, J.) e55539. ISSN: 2050-084X. doi:10.7554/eLife.55539 (July 28, 2020).

15. Goldshtein, A. et al. Onboard Recordings Reveal How Bats Maneuver under Severe Acoustic Interference. Proceedings of the National Academy of Sciences 122, e2407810122. doi:10.1073/pnas.2407810122 (Apr. 8, 2025).

16. Moss, C. F., Bohn, K., Gilkenson, H. & Surlykke, A. Active Listening for Spatial Orientation in a Complex Auditory Scene. PLOS Biology 4, e79. ISSN: 1545-7885. doi:10.1371/journal.pbio.0040079 (Mar. 7, 2006).

17. Camhi, J. M. Neuroethology: Nerve Cells and the Natural Behaviour of Animals. 416 pp. (Sinauer Associates, 1984).

18. Vanderelst, D. & Peremans, H. How Swarming Bats Can Use the Collective Soundscape for Obstacle Avoidance. PLOS Computational Biology 21, e1013013. ISSN: 1553-7358. doi:10.1371/journal.pcbi.1013013 (May 15, 2025).

19. Umadi, R. & Firzlaff, U. Biosonar Responsivity Sets the Stage for the Terminal Buzz https://www.biorxiv.org/content/10.1101/2025.06.16.659925v2 (2025). Pre-published.

20. Umadi, R. Temporal Feasibility Constraints on Wingbeat-Call Synchrony in Actively Echolocating Bats https://www.biorxiv.org/content/10.1101/2025.06.18.660328v2 (2025). Pre-published.

21. Couzin, I. D., Krause, J., Franks, N. R. & Levin, S. A. Effective Leadership and Decision-Making in Animal Groups on the Move. Nature 433, 513–516. ISSN: 1476-4687. doi:10.1038/nature03236 (Feb. 2005).

22. Middleton, D. An Introduction to Statistical Communication Theory 1170 pp. (New York, McGraw-Hill, 1960).

23. Helstrom, C. W. Statistical Theory of Signal Detection: Second Edition, Revised And… (1968).

24. Baddeley, A., Rubak, E. & Turner, R. Spatial Point Patterns: Methodology and Applications with R 828 pp. ISBN: 978-0-429-16170-4. doi:10.1201/b19708 (Chapman and Hall/CRC, New York, Nov. 11, 2015).

25. Umadi, R. Echolocating Bat Swarm Simulation version 1.0. July 2025. doi:10.6084/m9.figshare.29482268.v1.

26. Ahlswede, R. General Theory of Information Transfer: Updated. Discrete Applied Mathematics. General Theory of Information Transfer and Combinatorics 156, 1348–1388. ISSN: 0166-218X. doi:10.1016/j.dam.2007.07.007 (May 1, 2008).

27. Wilkinson, M. G. T., Wang, X. (, Cowan, N. J. & Moss, C. F. Echolocating Bats Adjust Sonar Call Features and Head/Ear Position as They Track Moving Targets in the Presence of Clutter. The Journal of the Acoustical Society of America 157, 2236–2247. ISSN: 0001-4966. doi:10.1121/10.0036252 (Mar. 27, 2025).

28. Lin, Y., Abaid, N. & Müller, R. Bats Adjust Their Pulse Emission Rates with Swarm Size in the Field. The Journal of the Acoustical Society of America 140, 4318. ISSN: 1520-8524. doi:10.1121/1.4971331. PMID: 28040047 (Dec. 2016).

29. Habersetzer, J. Adaptive Echolocation Sounds in the Bat Rhinopoma Hardwickei: A Field Study. Journal of Comparative Physiology ? A 144, 559–566. ISSN: 0340-7594, 1432-1351. doi:10.1007/BF01326841 (Dec. 1981).

30. Gillam, E. H., Hristov, N. I., Kunz, T. H. & McCracken, G. F. Echolocation Behavior of Brazilian Free-Tailed Bats during Dense Emergence Flights. Journal of Mammalogy 91, 967–975. ISSN: 0022-2372, 1545-1542. doi:10.1644/09-MAMM-A-302.1 (Aug. 16, 2010).

31. Chiu, C., Reddy, P. V., Xian, W., Krishnaprasad, P. S. & Moss, C. F. Effects of Competitive Prey Capture on Flight Behavior and Sonar Beam Pattern in Paired Big Brown Bats, Eptesicus Fuscus. Journal of Experimental Biology 213, 3348–3356. ISSN: 0022-0949. doi:10.1242/jeb.044818 (Oct. 1, 2010).

32. Wohlgemuth, M. & Moss, C. Active Listening in a Complex Environment. Proceedings of Meetings on Acoustics 19, 010030. ISSN: 1939-800X. doi:10.1121/1.4800959 (May 14, 2013).

33. Tuninetti, A., Ming, C., Hom, K. N., Simmons, J. A. & Simmons, A. M. Spatiotemporal Patterning of Acoustic Gaze in Echolocating Bats Navigating Gaps in Clutter. iScience 24, 102353. ISSN: 2589-0042. doi:10.1016/j.isci.2021.102353 (Apr. 23, 2021).

34. Weesner, A., Bentley, I., Fullerton, J. & Kloepper, L. Interaction Rules Guiding Collective Behaviour in Echolocating Bats. Animal Behaviour 206, 91–98. ISSN: 00033472. doi:10.1016/j.anbehav.2023.09.009 (Dec. 2023).

35. Salles, A., Diebold, C. A. & F. Moss, C. Bat Target Tracking Strategies for Prey Interception. Communicative & Integrative Biology 14, 37–40. ISSN: 1942-0889. doi:10.1080/19420889.2021.1898751 (Jan. 1, 2021).

36. Moss, C. F. & Surlykke, A. Auditory Scene Analysis by Echolocation in Bats. The Journal of the Acoustical Society of America 110, 2207–2226. ISSN: 0001-4966. doi:10.1121/1.1398051. PMID: 11681397 (2001).

37. Yang, X.-S. in Nature Inspired Cooperative Strategies for Optimization (NICSO 2010) (eds González, J. R., Pelta, D. A., Cruz, C., Terrazas, G. & Krasnogor, N.) 65–74 (Springer, Berlin, Heidelberg, 2010). ISBN: 978-3-642-12538-6. doi:10.1007/978-3-642-12538-6_6.

38. Trianni, V. & Campo, A. in (2015). ISBN: 978-3-662-43504-5. doi:10.1007/978-3-662-43505-2_71.

39. Müller, R. Bioinspiration from Bats and New Paradigms for Autonomy in Natural Environments. Bioinspiration & Biomimetics 19, 033001. ISSN: 1748-3190. doi:10.1088/1748-3190/ad311e (Apr. 2024).

40. Diebold, C. A., Salles, A. & Moss, C. F. Adaptive Echolocation and Flight Behaviors in Bats Can Inspire Technology Innovations for Sonar Tracking and Interception. Sensors 20, 2958. ISSN: 1424-8220. doi:10.3390/s20102958 (10 Jan. 2020).

41. Heilbron, M. & Chait, M. Great Expectations: Is There Evidence for Predictive Coding in Auditory Cortex? Neuroscience 389, 54–73. ISSN: 1873-7544. doi:10.1016/j.neuroscience.2017.07.061. PMID: 28782642 (Oct. 1, 2018).

42. Saberi, K. & Hickok, G. Forward Entrainment: Psychophysics, Neural Correlates, and Function. Psychonomic Bulletin & Review 30, 803–821. ISSN: 1531-5320. doi:10.3758/s13423-022-02220-y. PMID: 36460893 (June 2023).

